# The cAMP effector PKA mediates Moody GPCR signaling in *Drosophila* blood-brain barrier formation and maturation

**DOI:** 10.1101/2021.03.10.434865

**Authors:** Xiaoling Li, Richard Fetter, Tina Schwabe, Christophe Jung, Hermann Steller, Ulrike Gaul

## Abstract

The blood-brain barrier (BBB) of *Drosophila* is comprised of a thin epithelial layer of subperineural glia (SPG), which ensheath the nerve cord and insulate it against the potassium-rich hemolymph by forming intercellular septate junctions (SJs). Previously, we identified a novel Gi/Go protein-coupled receptor (GPCR), Moody, as a key factor in BBB formation at the embryonic stage. However, the molecular and cellular mechanisms of Moody signaling in BBB formation and maturation remain unclear. Here, we identify cAMP-dependent protein kinase A (PKA) as a crucial antagonistic Moody effector that is required for the formation, as well as for the continued SPG growth and BBB maintenance in the larva and adult stage. We show that PKA is enriched at the basal side of the SPG cell and that this polarized Moody/PKA pathway finely tunes the enormous cell growth and BBB integrity, by precisely regulating the actomyosin contractility, vesicle trafficking, and the proper SJ organization in a highly coordinated spatiotemporal manner. These effects are mediated in part by PKA’s molecular targets MLCK and Rho1. Moreover, 3D reconstruction of SJ ultrastructure demonstrates that the continuity of individual SJ segments and not their total length is crucial for generating a proper paracellular seal. Based on these findings, we propose a model that polarized Moody/PKA signaling plays a central role in controlling the cell growth and maintaining BBB integrity during the continuous morphogenesis of the SPG secondary epithelium, which is critical for maintain tissue size and brain homeostasis during organogenesis.

## Introduction

The blood-brain barrier (BBB) is a complex physical barrier between the nervous system and the peripheral circulatory system that regulate CNS homeostasis to ensure proper neuronal function. The *Drosophila* BBB is established by a thin epithelium of subperineural glia (SPG), which ensheath and insulate the nervous system against the potassium-rich hemolymph by forming intercellular septate junctions (SJs) (Bainton et al., 2005; Carlson et al., 2000; Edwards et al., 1993). The SPG epithelium is formed as a result of a mesenchymal-epithelial transition (MET), similar to other secondary epithelia such as heart and midgut. SPG cells only increase in number in embryogenesis but not in morphogenesis, and rather increase their size by polyploidization (Unhavaithaya and Orr-Weaver, 2012). Polyploidy in SPG is necessary to coordinate cell growth and BBB integrity either by Notch signaling or miR-285– Yki/Mask signaling during CNS development at the larval stage (Li et al., 2017; Unhavaithaya and Orr-Weaver, 2012; Von Stetina et al., 2018). SPG cells lack the apical markers present in primary epithelia (Crumbs, Bazooka), they have no contiguous zonula adherens and therefore rely on their SJ belt for epithelial cohesion and preventing paracellular diffusion and seal the BBB (Schwabe et al., 2005; Stork et al., 2008; Tepass, 2012; Tepass et al., 2001).

SJs are the crucial barrier junctions in invertebrates and functionally equivalent to vertebrate tight junctions; both junctions share Claudins as key components (Izumi and Furuse, 2014). Structurally and molecularly, SJs are homologous to the vertebrate paranodal junction (for review see (Banerjee et al., 2006; Salzer et al., 2008)). They consist of a core mutual interdependence protein complex, including transmembrane and cytoplasmic proteins, such as Neurexin-IV (Nrx-IV), Neuroglian (Nrg), the Na/K-ATPase (ATPα and Nrv2), the claudin Megatrachea (Mega), Sinous, Coracle (Cora), and the tetraspan Pasiflora protein family (Oshima and Fehon, 2011). In addition to the above-listed proteins, several GPI-anchored proteins, including Ly6-domain proteins Boudin, Crooked, Crimpled, and Coiled (Hijazi et al., 2011; Hijazi et al., 2009; Syed et al., 2011; Tempesta et al., 2017), Lachesin (Llimargas et al., 2004), Contactin (Faivre-Sarrailh et al., 2004), the tetraspan Pasiflora protein family(Deligiannaki et al., 2015) and Undicht (Petri et al., 2019), which are all found to be required for the SJ complex formation and proper membrane trafficking. The intracellular signaling pathways that control the assembly and maintenance of SJs are just beginning to be elucidated.

We have previously identified a novel GPCR signaling pathway that is required for the proper organization of SJ belts between neighboring SPG at the embryonic stage, consisting of the receptor Moody, two hetero-trimeric G proteins (Gαiβγ, Gαoβγ), and the RGS protein Loco. Both gain and loss of Moody signaling lead to non-synchronized growth of SPG cells, resulting in disorganized cell-contacts and shortened SJs and therefore, a leaky BBB (Schwabe et al., 2005; Schwabe et al., 2017). The phenotype of Moody is weaker than that of downstream pathway components including Loco and Gβ13F, suggesting that additional receptors provide input into the trimeric G protein signaling pathway. Gγ1 signaling has been shown to regulate the proper localization of SJ proteins in the embryonic heart (Yi et al., 2008). Despite its critical role in BBB formation, the underlying mechanisms connecting G protein signaling to continued SPG cell growth and the proper SJ organization during the development and maturation of BBB are still poorly understood.

One of the principal trimeric G protein effectors is Adenylate Cyclase (AC). AC is inhibited by the G proteins Gαi/Gαo and Gβγ, leading to decreased levels of the second messenger cAMP. The prime effector of cAMP, in turn, is cAMP-dependent protein kinase A (PKA), a serine/threonine kinase. PKA is inactive as a tetrameric holoenzyme, which consists of two identical catalytic and two regulatory subunits. Binding of cAMP to the regulatory units releases and activates the catalytic subunits (Taylor et al., 1990). PKA transmits the signal to downstream effectors by phosphorylating multiple substrates which participate in many different processes, from signal transduction to regulation of cell shape and ion channel conductivity (Shabb, 2001). In *Drosophila*, PKA has been studied as a component of GPCR signaling in the Hedgehog pathway during development (Li et al., 1995; Marks and Kalderon, 2011), and in neurotransmitter receptor pathways during learning and memory (Chen and Ganetzky, 2012; Guan et al., 2011; Li et al., 1996; Renger et al., 2000). PKA also regulates microtubule organization and mRNA localization during oogenesis (Lane and Kalderon, 1993, 1994, 1995). In vertebrates, cAMP/PKA signaling is known to play a central role within different subcellular regions, including the regulation of actomyosin contractility and localized cell protrusion in directional cell migration (Howe, 2004; Lim et al., 2008; Tkachenko et al., 2011); intracellular membrane trafficking (exocytosis, endocytosis and transcytosis) in relation to the dynamics of epithelial surface domains in developmental processes and organ function (Wojtal et al., 2008); and the regulation of endothelial tight junction (TJ) with diverse actions and uncleard mechanisms in different endothelial cells models(Cong and Kong, 2020).

Here, we report results from a comprehensive *in vivo* analysis of the molecular and cellular mechanisms of Moody signaling in the SPG. We show that PKA is a key downstream effector responsible for the salient phenotypic outcomes, and that it acts by modulating actomyosin contractility via MLCK and Rho1. The strong phenotypic effects of PKA gain- and loss-of-function permit a detailed dissection of the organization of cell-cell contacts as driven by Moody/PKA signaling and allow us to track its role in the continued growth of the SPG during larval stages. We observe asymmetric and opposing subcellular distributions of Moody and PKA, providing novel insight into the establishment of apical-basal polarity in the SPG as a secondary epithelium, as well as its morphogenetic function. We present a 3D reconstruction of SJ ultrastructure using serial section Transmission Electron Microscopy (ssTEM) under different PKA activity levels. This new analysis reveals a strict coupling of total cell contact and SJ areas, but also suggests that it is the continuity of individual SJ segments and not total SJ width that is essential for normal BBB insulation. Altogether, our data reveal a previously unrecognized role of GPCR/PKA in maintaining enormous SPG cell growth and it’s sealing capability by regulating actomyosin contractility and the proper SJ organization in BBB formation and maturation, which touches the fundamental aspects of remodeling cytoskeletal network spatiotemporally - a common processes but with different mechanisms in morphogenesis.

## Results

### PKA is required for Moody-regulated BBB formation

To identify molecules that act downstream of Moody signaling in BBB formation, we examined genes known to be involved in GPCR signaling, such as PkaC1, PI3K, PTEN, PLC, and Rap1. We tested BBB permeability in genomic mutants or transgenic RNAi knockdowns of these GPCR effectors, by injecting a fluorescent dye into the body cavity and determining its penetration into the CNS using confocal imaging. We found that zygotic mutants of the PKA catalytic subunit PkaC1 (originally named DC0 in *Drosophila*), namely the two null alleles *PkaC1*^*B3*^ and *PkaC1*^*H2*^ as well as the hypomorphic allele *PkaC1*^*A13*^ (Kalderon and Rubin, 1988) show severe CNS insulation defects (Figure 1A-B), similar in strength to zygotic mutants of the negative regulator *loco*. By contrast, the removal of the other candidates had no effect (data not shown). *PkaC1* has both maternal and zygotic components, and its maternal contribution perdures until late embryogenesis (Lane and Kalderon, 1993). The BBB defect we observe could explain the morphologically inconspicuous embryonic lethality of *PkaC1* zygotic null mutants (Lane and Kalderon, 1993). To rule out the possibility that the observed BBB defects are caused by glial cell fate or migration defects, we examined the presence and position of SPG using an antibody against the pan-glial, nuclear protein Reversed polarity (Repo) (Halter et al., 1995). In *PkaC1* zygotic mutants, the full set of SPG is present on the surface of the nerve cord, although the position of the nuclei is more variable than in WT (Figure 1C), an effect that is also observed in known mutants of the Moody signaling pathway (Granderath et al., 1999; Schwabe et al., 2005).

Since SJs are the principal structure providing BBB insulation and are disrupted in Moody pathway mutants (Schwabe et al., 2005; Schwabe et al., 2017), we sought to characterize the SJ morphology in PkaC1 mutants. We performed ultrastructural analysis of SJs in late embryos (AEL 22-23h) by Transmission Electron Microscopy (TEM) using high pressure freezing fixation. In WT, the SJs are extended, well-organized structures that retain orientation in the same plane over long distances (Figure 1D). In contrast, in *PkaC1*^*H2*^ zygotic mutants, the overall organization of SJs appears perturbed, and their length, as measured in random single sections, is significantly shorter than in WT (Figure 1D-E); very similar phenotypic defects are observed in *moody* and *loco* zygotic mutants (Schwabe et al., 2005).

**Figure 1.**
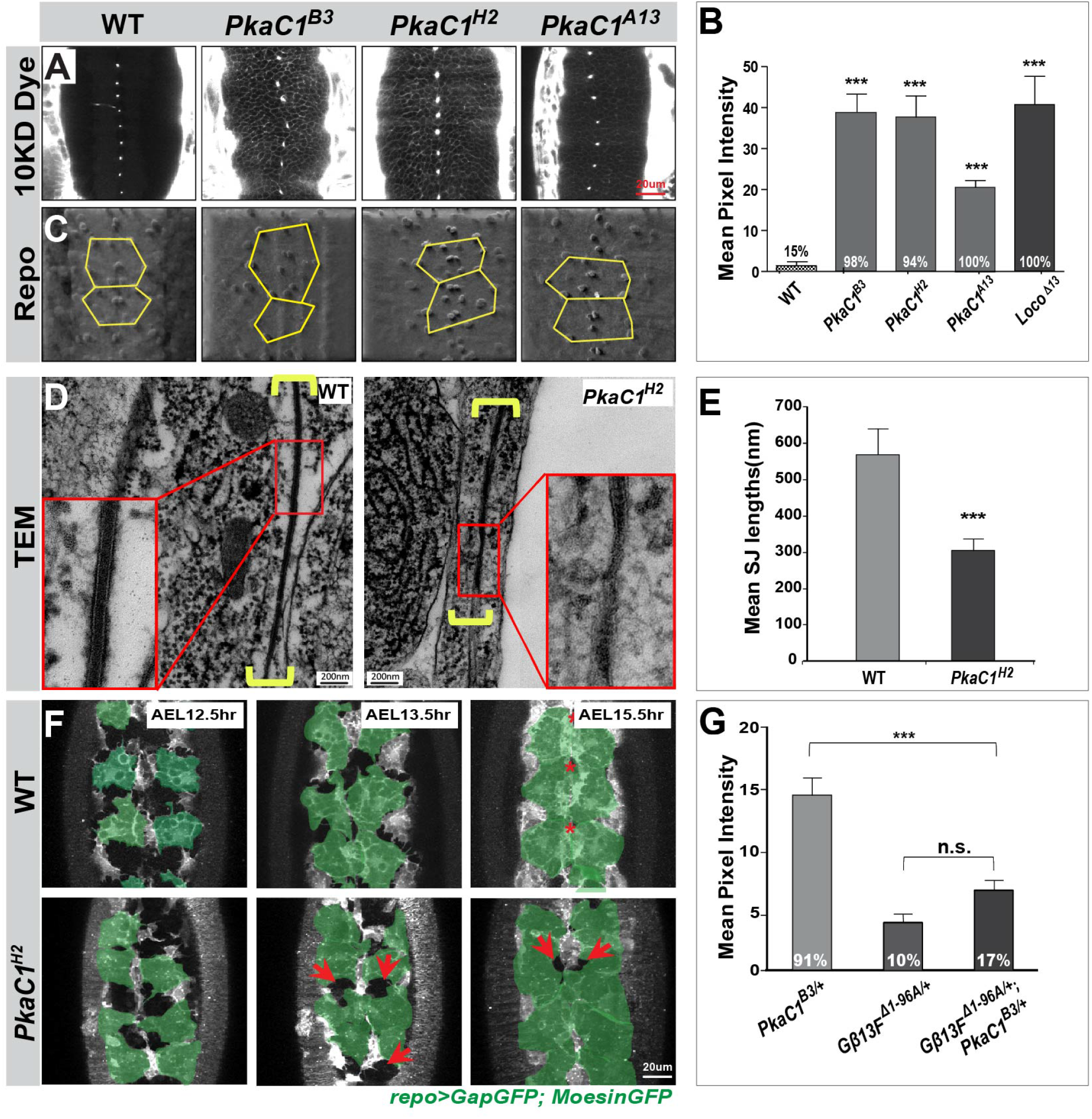
PKA is required for BBB formation and acts in the Moody signaling pathway. (A) Single confocal sections of dye-injected embryos of WT and PKA zygotic mutants. (B) Quantification of the dye penetration assay. Columns represent the intensity of dye penetration into the nerve cord as measured by the mean pixel intensity (see Experimental Procedures), ±SEM, n=16-41. *Loco*^*Δ13*^ zygotic mutants serve as positive controls. (C) Repo staining revealing the number and positions of SPG nuclei in WT and PKA zygotic mutants using an illuminated projection to highlight the ventral surface of the nerve cord. (D) Transmission electron micrographs of the interface of neighboring SPG in late WT and *PkaC1*^*H2*^ zygotic mutant embryos. Yellow brackets delineate the SJ ultrastructure; high magnifications are shown in red boxes. (E) Quantification of SJ length in WT and *PkaC1*^*H2*^ mutants (see Experimental Procedures). Columns represent mean SJ length as measured in random nerve cord sections, ±SEM, n=56-70. (F) Time-lapse recording of BBB closure in embryos of WT and *PKA* zygotic mutants. 6 µm confocal stacks are shown; in each image, 4-6 ventral SPG are highlighted (green); midline channels (stars) and retarded growth (arrows) are marked. (G) Dominant genetic interactions between *PkaC1*^*B3*^ and *Gβ13F*^*Δ1-96A*^ as quantified by dye penetration in the embryo. Columns represent the intensity of dye penetration as measured by the mean pixel intensity, ±SEM, n=34-48. In (B) and (G), the percentage of embryos showing the dye penetration is indicated at the bottom of each column. Brackets and asterisks in (B), (E) and (G) indicate statistical significance levels as assessed by one-way ANOVA with Dunnett’s multiple comparisons test (B) and (G) or the two-tailed Student’s t-test (E), n.s. p> 0.05; *p<0.05; **p<0.01; ***p<0.001.

To explore the role of PkaC1 during development of the BBB, we performed time-lapse recordings of SPG epithelium formation. The SPG arise in the ventro-lateral neuroectoderm and migrate to the surface of the developing nerve cord (Ito et al., 1995), where they spread until they reach their neighbors and form intercellular SJs (Schwabe et al., 2005; Schwabe et al., 2017). To monitor the changes in SPG morphology during the closure process, we expressed the membrane marker *GapGFP* and the actin marker *MoesinGFP* using the pan-glial driver *repoGAL4* (Schwabe et al., 2017) (Figure 1F, movies S1 and S2). In WT embryos, SPG are relatively uniform in cell size and shape, and grow to form cell-cell contacts in a highly synchronized manner. By 15.5 h of development, the glial sheet is closed (Figure 1F). By contrast, SPG in *PkaC1*^*H2*^ zygotic mutants show increased variability in size and shape, and their spreading and contact formation is less well coordinated. This results in patchy cell-cell contacts with gaps of variable sizes (Figure 1F). Moreover, the complete closure of the SPG epithelium is delayed compared to WT (Figure 1F). Again, the defects observed in PKA loss-of-function are similar to those in Moody pathway mutants (Schwabe et al., 2017).

Our results show that *PkaC1* is required for BBB integrity, proper SJ organization, and SPG epithelium formation, in all cases closely mimicking the phenotypes observed for known Moody signaling components. Given these similarities, we sought to determine whether PKA participates in the Moody pathway by performing dominant genetic interaction experiments. Notably, we found that embryos heterozygous for *PkaC1* null alleles, which are known to have ∼50% of wild type PkaC1 activity, show mild BBB permeability defects (Figure 1G). Therefore, we used *PkaC1*^*B3*^ heterozygous mutants as a sensitized genetic background and removed one genomic copy of different Moody pathway components, including Moody, Loco, Gαo, Gαi, and Gβ13F (Schwabe et al., 2005), to determine whether any synergistic or antagonistic interactions are observed. We found that the dye penetration defects of *PkaC1* heterozygous mutants are significantly reduced by removing one genomic copy of *Gβ13F* or *loco* (Figure 1G and S1); removal of one genomic copy of *Gβ13F* or *loco* on their own have no effect. These genetic interactions indicate that PkaC1 is indeed part of the Moody signaling pathway. Removal of single copies of other pathway components showed either a mild, non-significant or no effect in a *PkaC1*^*B3*^ background, suggesting that they are less dosage-sensitive (Figure S1).

### PKA is required for BBB continued growth in larvae and BBB maintenance in adults

For a more detailed analysis of PkaC1 function in BBB regulation, we turned to the SPG epithelium in third instar larvae. During the larval stage, no additional SPG cells are generated, instead the existing SPG cells grow enormously in size to maintain integrity of the BBB (Li et al., 2017; Unhavaithaya and Orr-Weaver, 2012). By third instar, they have roughly doubled in size and are accessible via dissection of the CNS, which greatly facilitates the microscopic analysis. PKA activity in larvae can be manipulated specifically using the SPG-specific driver *moodyGAL4* (Bainton et al., 2005; Schwabe et al., 2005), which becomes active only after epithelial closure and BBB sealing are completed in stage 17 embryos. PKA can be reduced by expression of transgenic RNAi targeting the PKA catalytic subunit C1 (*moody>PkaC1-RNAi*). On the other hand PKA can be elevated by expression of a mouse constitutively active PKA catalytic subunit (*moody*>*mPkaC1*)*(Zhou et al., 2006). We first examined whether normal Moody/PKA activity is required for BBB integrity during larval stages. To address this question, we developed a dye penetration assay to measure BBB permeability in cephalic complexes of third instar larval. This assay is similar to the one we performed in the late embryo, but with some important modifications (for details see Experimental Procedures). Interestingly, both elevated and reduced activity of Moody (*moody>LocoRNAi* and *moody>moodyRNAi*) and PKA (*moody>mPKAC1\** and *moody>PKAC1RNAi*) in SPG resulted in severe BBB insulation defects (Figure 2A and 2D). This strongly suggests that Moody/PKA signaling plays a crucial role in the continued growth of the BBB during larval stages. These effects were not merely carried over from the embryo, since under *moody* driver caused only mild dye penetration defects in embryos (Figure S2). Given that Moody activity has been implicated in the maintenance of the BBB in the adult (Bainton et al., 2005), we also sought to knockdown PKA specifically in the adult SPG (*tubGal80ts, moody>PkaC1RNAi*) and measure the resulting effects (see Experimental Procedures). We indeed observed the dye penetrated the blood-eye barrier (Figure 2F), indicating that PKA is also required for BBB integrity function in the adult.

**Figure 2.**
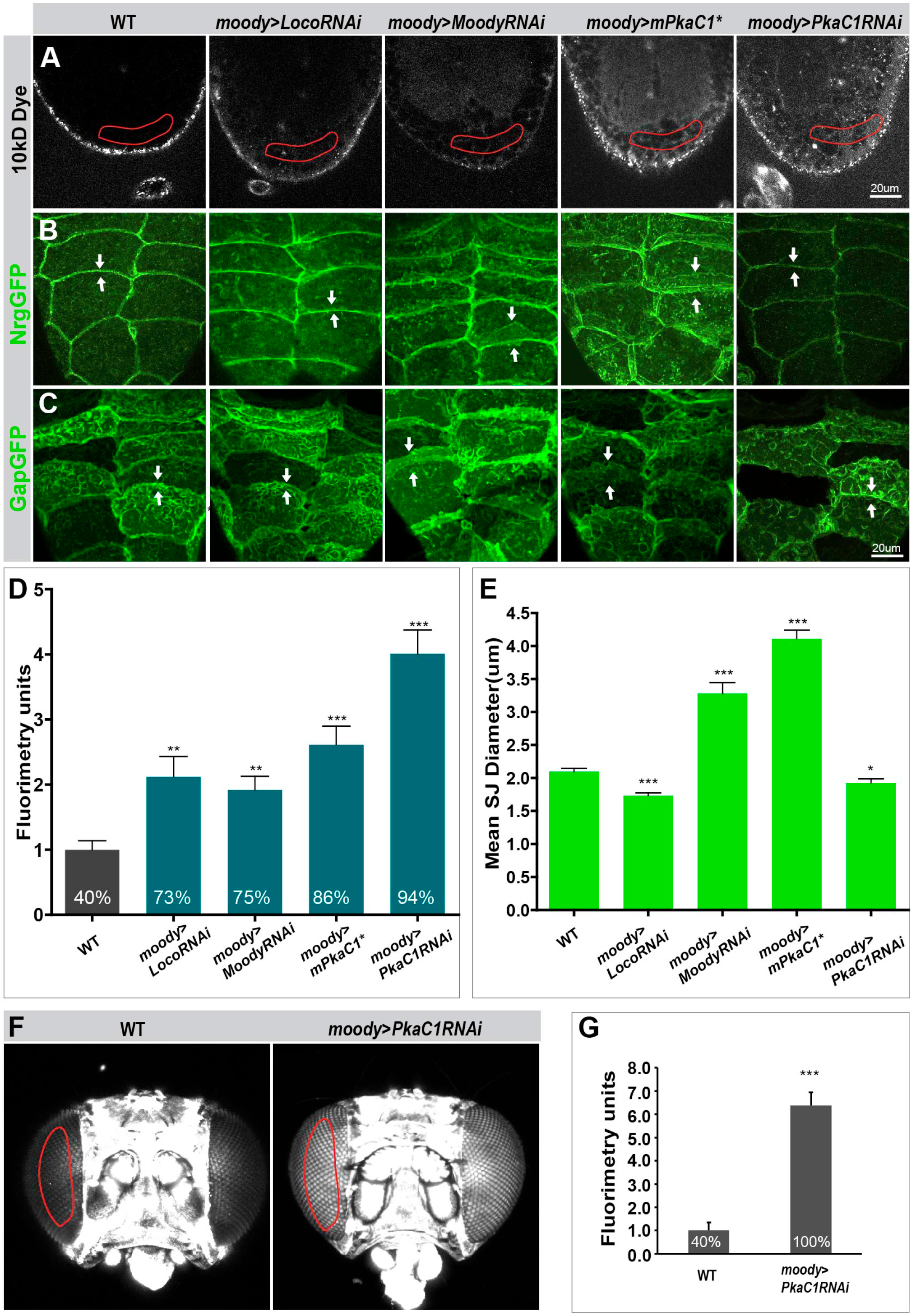
Moody/PKA signaling is required for BBB growth in the larva and for BBB maintenance in the adult. (A) Single confocal sections of dye injected third instar larval nerve cords under different Moody/PKA activity levels. (B-C) Morphology of SPG SJ belts and membrane overlap at different Moody/PKA activity levels, as visualized by SJ markers NrgGFP (B), and the membrane marker GapGFP (C). (D) Quantification of the dye penetration assay. Columns represent intensity of dye penetration as measured by mean pixel intensity (see Experimental Procedures), ±SEM, n=44-88. The percentage of larva showing dye penetration is indicated at the bottom of each column. (E) Quantification of the diameter of SJ belts under different GPCR/PKA activity levels, using the SJ marker NrgGFP. ±SEM, n=7-28. (F) Dye penetration in adult flies as shown in z-projections of dye-injected adult heads. (G) Quantification of dye penetration in adult eye. Columns represent intensity of dye penetration as measured by mean pixel intensity in adult eye (see Experimental Procedures), ±SEM, n=30 and 18. Asterisks in (D), (E), and (G) indicate significance levels of comparisons based on Welch’s ANOVA with Dunnett’s T3 multiple comparisons test (D) and (E) or the two-tailed Student’s t-test (G), n.s. p>0.05; *p<0.05; **p<0.01, ***p<0.001.

In order to better understand the cause of BBB permeability under conditions where Moody/PKA is changed, we examined SJ morphology in larvae. Most core SJ components show interdependence for correct localization and barrier function, with removal of one component sufficient to abolish SJ function (Behr et al., 2003; Genova and Fehon, 2003; Hijazi et al., 2011; Oshima and Fehon, 2011; Wu et al., 2004). We therefore asked whether PKA activity levels affect the distribution of different SJ components. Using both live imaging (*NrgGFP, LacGFP, NrxIVGFP*) and immunohistochemistry (Mega), we found that the circumferential SJ belts and outlines of SPG were marked nicely in WT (Figure 2C and S3). Strikingly, upon either reduction of Moody activity or elevated PKA activity, the SJ belt staining became much broader and more diffuse than in WT (Figure 2B). This suggests extensive plasma membrane overlap between neighboring SPG cells (Figure 2C). To confirm this idea, we introduced the membrane marker *gapGFP*, and indeed observed increased membrane overlap compared to WT (Figure 2C). Conversely, both elevated Moody activity and reduced PKA activity resulted in thinner SJ belts and reduced membrane (Figure 2B-C). To quantify these changes, we measured the mean width of the SJ belts under different PKA activity levels (Figure 2E; Experimental Procedures). The mean width of SJ belts increased with elevated PKA activity/reduced Moody activity and decreased under inverse conditions compared to WT (Figure 2H). These data demonstrate that Moody and PKA are required for the continued growth of the BBB and the proper organization of SJs during larval stages. Unlike the barrier defect, these morphological data reveal a monotonic relationship between PKA activity, membrane overlap and the amount of SJ components in the area of cell contact. The fact that the cellular defects of reduced Moody activity match those of elevated PKA activity, and vice versa, provides further evidence that PKA acts as an antagonistic effector of Moody signaling.

### PKA regulates the cytoskeleton and vesicle traffic in SPG

We had previously reported that the Moody pathway regulates the organization of cortical actin and thus the cell shape of SPG during late embryogenesis (Schwabe et al., 2005; Schwabe et al., 2017). Moreover, we proposed, based on the developmental timeline, that this in turn affects the positioning of SJ material along the lateral membrane. Given that the most striking phenotype caused by altered PKA activity is the extent of membrane overlap, we sought to further explore if PKA functions by regulating the cytoskeleton in SPG.

For this purpose, we examined the intracellular distribution of the actin cytoskeleton in the SPG at different PKA levels. As live markers we used *GFPactin*, which labels the entire actin cytoskeleton, *RFPmoesin* (Schwabe et al., 2005), which preferentially labels the cortical actin, the presumptive general MT marker *TauGFP* (Jarecki et al., 1999), the plus-end marker *EB1GFP* (Rogers et al., 2004), and the minus-end marker *NodGFP* (Clark et al., 1997; Cui et al., 2005) (Figure 3A-D, and data not shown). In response to changes in PKA activity, all markers showed altered distributions similar to those observed with SJ markers. Specifically, elevated PKA activity caused all markers to become enriched at the cell cortex, consistent with the broader membrane overlap between neighboring SPG (Figure 3A-D, middle column). Conversely, upon reducing PKA activity, all markers were reduced or depleted from the cell cortex, consistent with reduced contact area between SPG (Figure 3A-D, right column). Thus, PKA signaling profoundly reorganizes the actin and MT cytoskeleton, thereby regulating the membrane overlap formed between neighboring SPG.

**Figure 3.**
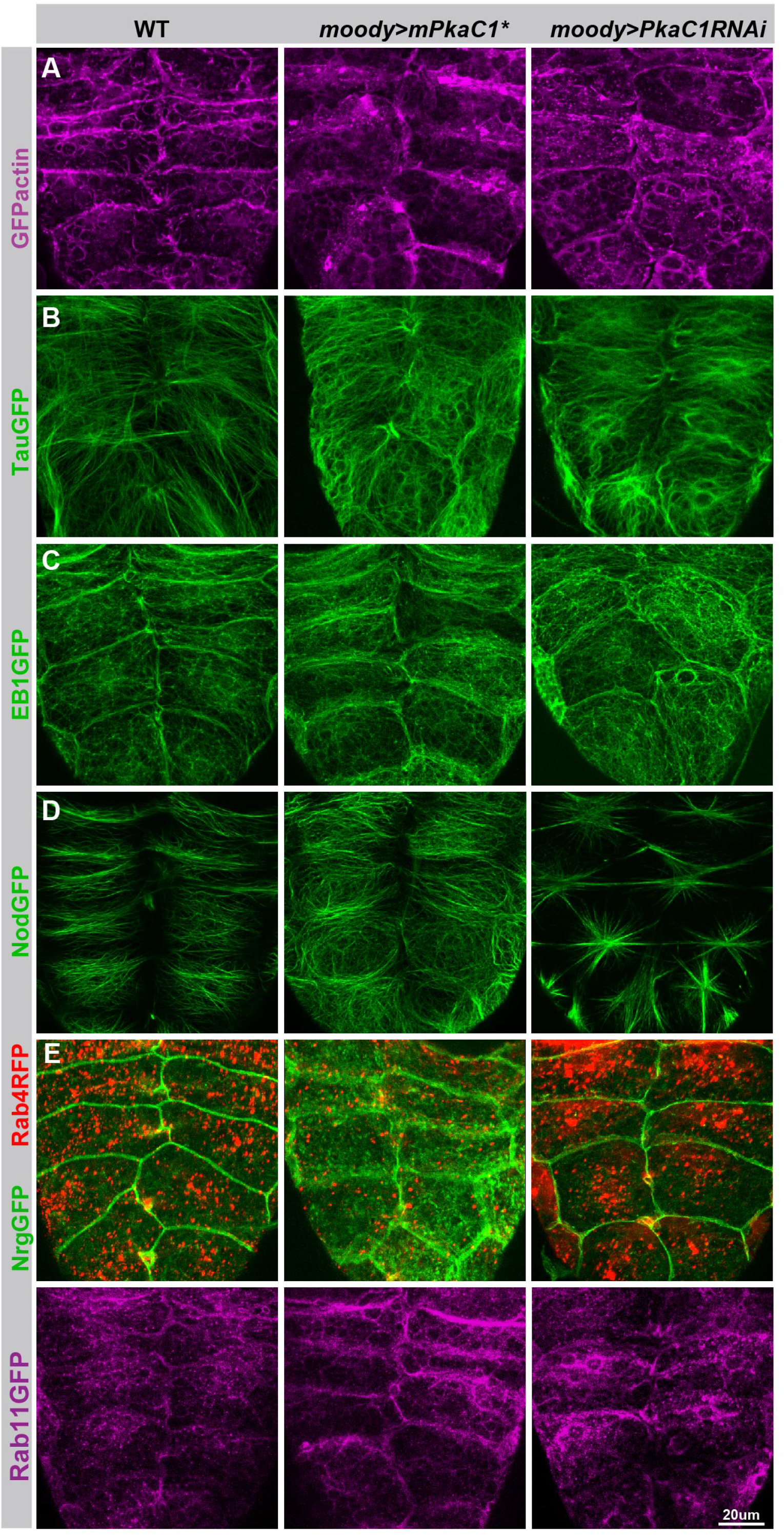
PKA regulates the cytoskeleton and vesicle distribution in SPG. Under different PKA activity levels, the actin cytoskeleton is visualized by GFPactin (A), the microtubule cytoskeleton by the general MT marker TauGFP (B), the plus-end marker EB1GFP (C) and the minus-end marker NodGFP (D), the cellular distribution of vesicles by the early endosome markers Rab4RFP (E) and Rab11GFP (F) with the SJ marker NrgGFP labeling the cell periphery of the SPG (E).

Since PKA has been shown to affect vesicle trafficking in epithelial cells and neurons (Renger et al., 2000; Vasin et al., 2014; Wojtal et al., 2008; Zhang et al., 2007), we investigated if PKA signaling has a similar role during continued SPG cell growth. We introduced two live markers, *Rab4RFP*, which labels all the early endosomes (Figure 3E), and *Rab11GFP* (Artiushin et al., 2018), which labels both early and recycling endosomes (Figure 3F). We observe significant changes in the cellular distribution of vesicle populations. Specifically, Rab4- and Rab11-labeled endosomes were differentially enriched in the cell periphery when PKA activity is increased, and surrounded the nucleus when PKA was reduced, as compared to their broader cytoplasmic distribution profile in WT (Figure 3E and 3F). Therefore, upon increasing levels of PKA all cytoskeletal and vesicular markers responded with monotonic changes, resulting in their increasing accumulation at the cell cortex of SPG.

### The continuity of SJ belt is essential for BBB function as revealed by ssTEM

While PKA gain- and loss-of-function show opposite morphologies of membrane overlap and SJ belt by light microscopy, they both result in a compromised leaky BBB. To better understand this incongruence, we sought to analyze membrane morphology at a higher resolution. Due to the small size of SJs (20-30nm), structural aspects can be analyzed conclusively only by electron microscopy. In the past, the acquisition and analysis of a complete series of TEM sections required an enormous effort; as a consequence, studies of SJ structure have mostly been restricted to random sections (Carlson et al., 2000; Hartenstein, 2011; Stork et al., 2008; Tepass and Hartenstein, 1994). The problem has now become solvable, using digital image recording (Suloway et al., 2005) and specialized software (Fiji, TrakEM2)(Cardona et al., 2012; Schindelin et al., 2012) for both image acquisition and post-processing. Therefore, we performed serial section TEM, followed by computer-aided reconstruction of TEM stacks to resolve the 3D ultrastructure of cell contacts and SJs under different PKA activity levels at third instar larva (Figure 4A and 4B). This is the first time that a contiguous SJ belt between neighboring SPG at nanometer resolution is presented.

**Figure 4.**
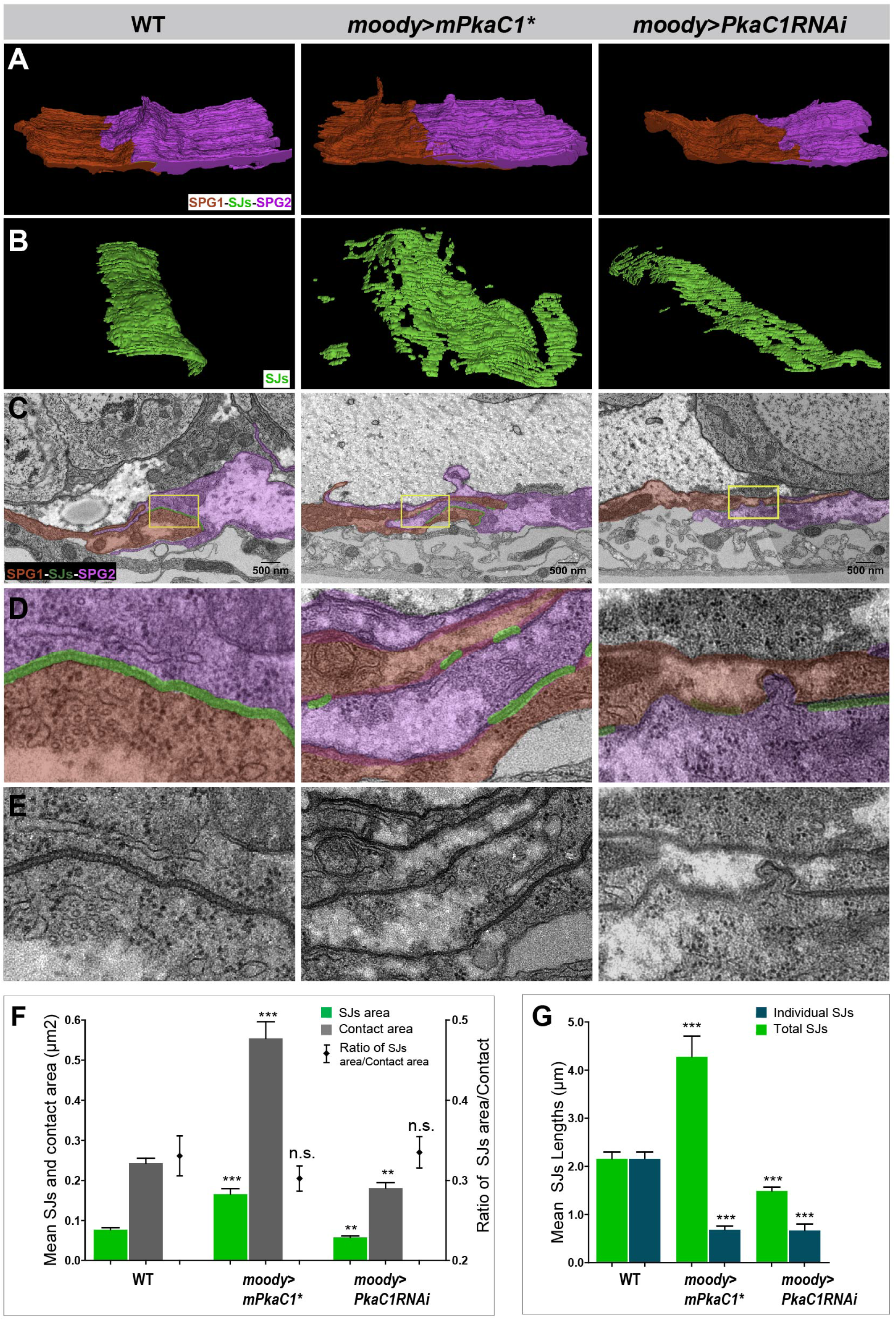
The continuity of the SJ belt is essential for BBB function as revealed by ssTEM. (A-E) SJ ultrastructure at the interface of neighboring SPG in third instar larvae under different PKA activity levels. SPG1, its neighbor SPG2 and their shared SJs are colored or shaded in red, magenta and green, respectively. (A and B) A 3D model of SJ ultrastructure generated by ssTEM. (C) Representative sections of SJs. (D and E) High magnification views of boxed regions in C with and without shading. (F) Quantification of SJ surface area (green column) and the contact area (grey column), and the ratio between the two (black point) under different PKA activity levels, ±SEM, n=15-21. (G) Quantification of the mean length of individual SJ segments (green) and the mean total length of SJs (blue) under different PKA activity levels, measured in random nerve cord sections, ±SEM, n=9-92. Asterisks in (F-G) indicate significance levels of comparisons based on Welch’s ANOVA with Dunnett’s T3 multiple comparisons test, n.s. p>0.05; *p<0.05; **p<0.01, ***p<0.001.

In WT, the area of cell-cell contact is compact and well-defined, with a dense SJ belt covering ∼30% of the cell contact area (Figure 4A, 4B, and 4F). Upon elevated PKA activity, neighboring SPG show much deeper membrane overlap (Figure 4A-E). The areas of both cell contact and SJ coverage increase about two-fold compared with WT (Figure 4F), confirming the observations from confocal microscopy (Fig 3A-D), but the SJ belt is discontinuous and appears patchy (Figure 4B-E). This suggests that it is the continuity of the belt, rather than the total area covered by SJs, that is essential for generating the intercellular sealing capacity. To examine this question directly, we measured SJ length in randomly selected sections. Compared with WT, the mean length of individual SJ segments (0.69 ± 0.08 µm vs. 2.16 ± 0.14 µm, p<0.0001) is indeed significantly decreased, while the mean total length of SJs (4.28 ± 0.43 µm vs. 2.16 ± 0.14 µm, p= 0.000523) is significantly increased (Figure 4G).

Upon reducing PKA activity, the cell contacts and SJ area between neighboring SPGs were reduced, and the SJ belt became patchy as well (Figure 4A-E). In this case, both the mean total length of SJs (1.49 ± 0.08 µm vs. 2.16 ± 0.14 µm, p= 0.000878) and the mean length of individual SJ segments (0.67 ± 0.14 µm vs. 2.16 ± 0.14 µm, p<0.0001) were significantly shorter than in WT (Figure 4G). Intriguingly, the ratio of total SJ area to cell contact area remains constant at about 30% under all PKA activity conditions, despite the variable interdigitations between contacting SPG (Figure 4F).

Finally, SPG send apical protrusions into the neural cortex (Figure S4). These protrusions are much longer (2.01 ± 0.01 µm vs. 1.47 ± 0.09 µm, p=0.000230) upon elevated PKA activity and shorter than in WT (0.68±0.09 µm vs. 1.47±0.09 µm, p<0.0001) upon reduced PKA activity, suggesting that PKA activity more generally controls membrane protrusions and extension(Figure S4).

Taken together, our ultrastructural analyses and new 3D models support the light microscopic findings, and they provide superior quantification of the relevant parameters. Importantly, cell contact and SJ area, as well as total SJ content are monotonically correlated with PKA activity, while individual SJ segment length is not. This suggests that the discontinuity of the SJ belt is the main cause for the observed BBB permeability defects.

### The Moody/PKA signaling pathway is polarized in SPG

The SPG are very thin cells, measuring around 0.2 µm along the apical-basal axis. In the embryo, the hemolymph-facing basal surface of the SPG is covered by a basal lamina (Fessler et al., 1994; Olofsson and Page, 2005; Tepass and Hartenstein, 1994), while during larval stages, the Perineurial Glia (PNG) form a second sheath directly on top of the SPG epithelium, which then serves as the basal contact for the SPG (Stork et al., 2008). Consistent with its chemoprotective function, the Mdr65 transporter localizes to the hemolymph-facing, basal surface of the SPG, while Moody localizes to the CNS-facing, apical surface (Mayer et al., 2009). The shallow lateral compartment contains the SJs, which not only seal the paracellular space but also act as a fence and prevent diffusion of transmembrane proteins across the lateral compartment. The apical localization of Moody protein is dependent on the presence of SJs (Schwabe et al., 2017).

To visualize the subcellular protein distributions along the apical-basal axis, we labeled them together with the SPG nuclei (*moody>nucCherry)*. We examined the subcellular distribution of PkaC1 by immunohistochemistry (anti-PKA catalytic subunit antibody, which only bind to the catalytic subunits of PKA dissociated from the regulatory subunits of PKA after cAMP activation, not binding to the inactive holoznzyme), and found that active PkaC1 is enriched on the basal side of the SPG, and thus the opposite of the apically localized Moody (Figure 5A). This result is intriguing given PKA’s antagonistic role in Moody signaling and suggests that pathway activity may affect the localization of pathway components. The subcellular distribution of PkaC1 was indeed altered when Moody is knocked down. PkaC1 lost its basal intracellular localization and appeared spread out throughout the cytoplasm (Figure 5B). This suggests that apical Moody signaling is necessary for repressing apical PkaC1 protein accumulation, and that this polarized subcellular localization results from the antagonistic relationship between Moody and PKA.

**Figure 5.**
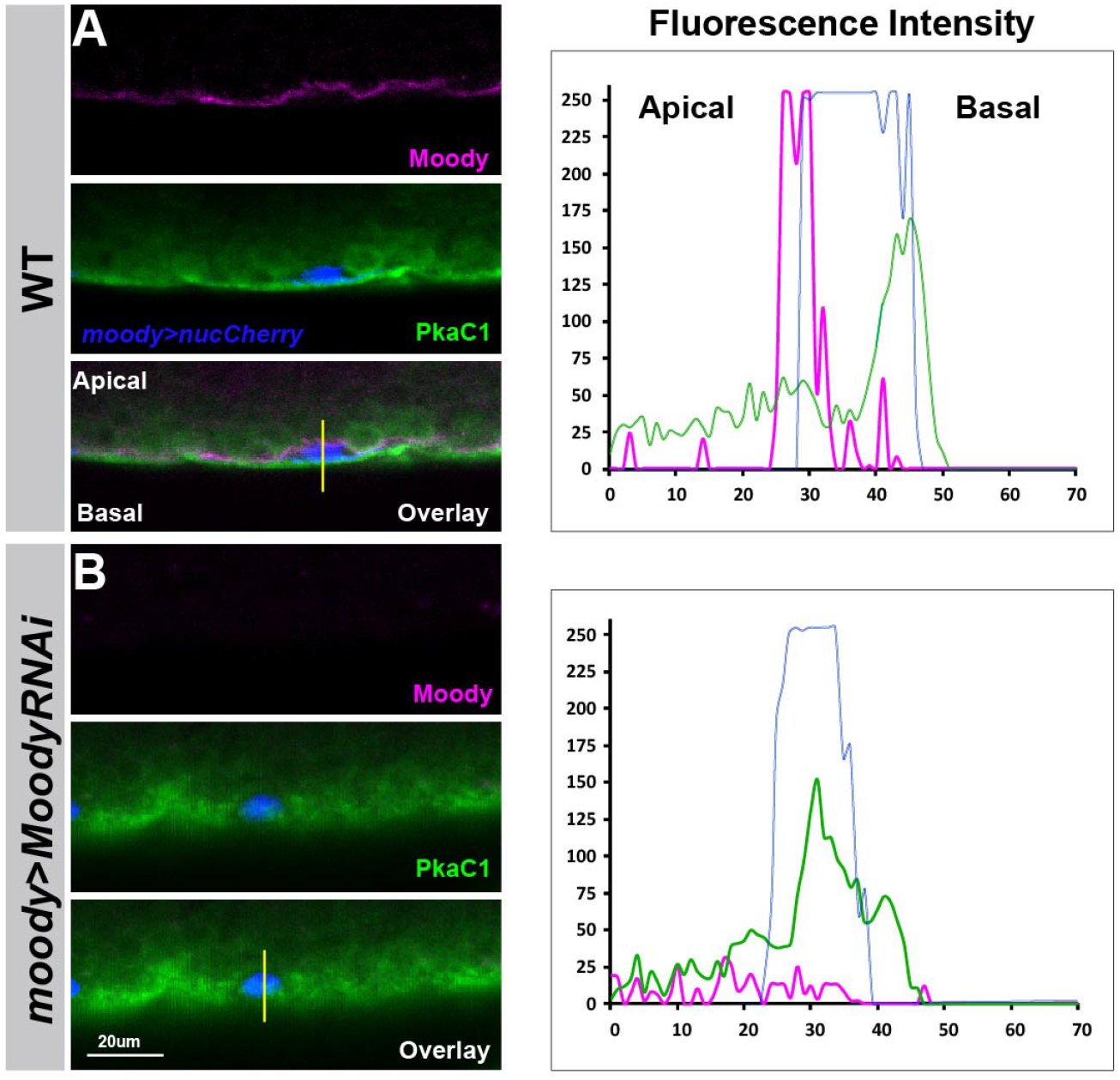
The Moody/PKA signaling pathway is polarized in SPG. The subcellular localization of the PKA catalytic subunit PkaC1 and Moody in SPG of third instar larvae in WT (A) and *moody* knockdown (B). Antibody labeling of Moody (magenta), of *Drosophila* PkaC1 or mouse PkaC1 (green), and of SPG nuclei (*moody>nucCherry*; red). Lateral views of the CNS/hemolymph border, with CNS facing top. On the right, line scans of fluorescence intensities for each channel along the apical-basal axis at the positions indicated.

### MLCK and Rho1 function as PKA targets in the SPG

Considering that the most pronounced effect of increasing PKA levels in the SPG is a commensurate increase in membrane overlap at the basolateral side, we sought to genetically identify PKA targets involved in this process. PKA is known to regulate actomyosin contractility by phosphorylating and inhibiting myosin light chain kinase (MLCK), which leads to a decrease in Myosin light chain (MLC) phosphorylation and a concomitant reduction of actomyosin contractility in cell migration and endothelial barrier (Garcia et al., 1995; Garcia et al., 1997; Howe, 2004; Tang et al., 2019; Verin et al., 1998). To determine whether MLCK is required for BBB function, we examined two MLCK zygotic mutants, *MLCK*^*02860*^ and *MLCKC*^*234*^, and detected moderate BBB permeability in the late embryo (Figure 6A and S5), indicating that MLCK plays a role in CNS insulation. Next, we asked whether PKA and MLCK function in the same signaling pathway using dominant genetic interaction experiments. We found that the BBB permeability of *PkaC1*^*B3*^ heterozygous mutants could be rescued by removing one parental copy of *MLCK* (*MLCK*^*02860*^ or *MLCK*^*C234*^) (Figure 6B). This suggests that MLCK interacts with PkaC1 in the SPG. Finally, we examined BBB insulation and SJ defects of *MLCK* zygotic mutant larva (*MLCK*^*C234*^). *MLCK*^*C234*^ mutant larvae showed significant BBB permeability and a widened SJ belt (Figure 6C-F) compared to WT (Figure 2B), but the phenotypes were milder than those of PKA overactivity (Figure 2B).

**Figure 6.**
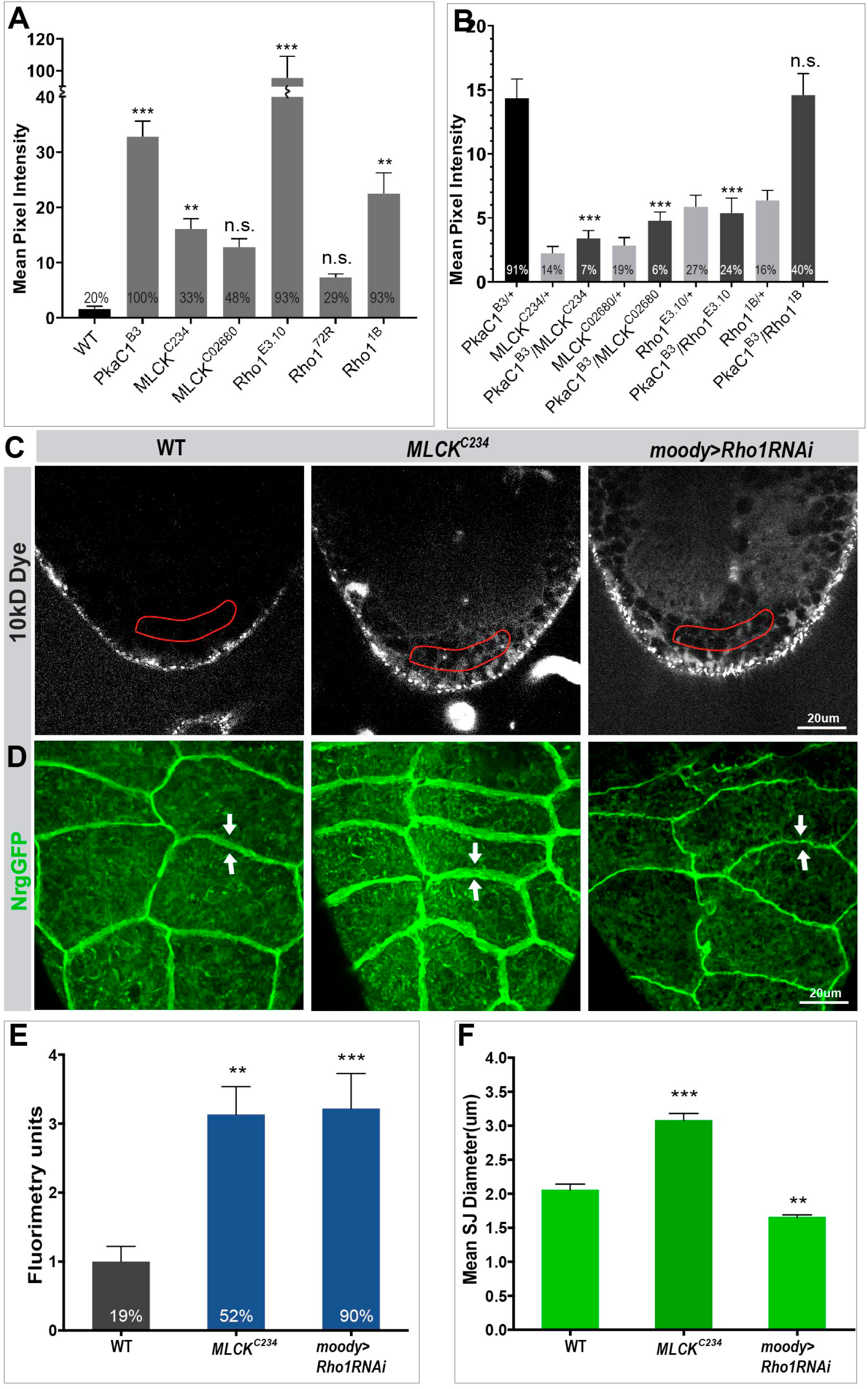
MLCK and Rho1 function as PKA targets in SPG. (A) Quantification of dye penetration effects in the embryo of *MLCK* and *Rho1*. (B) Dominant genetic interactions between *PkaC1*^*B3*^ and *MLCK* and *Rho1* mutant heterozygotes as quantified by dye penetration in the embryo. In (A) and (B), columns represent the strength of dye penetration into the nerve cord as measured by the mean pixel intensity, ±SEM, n=14-98. (C-D) BBB phenotype of *MLCK* zygotic mutant and SPG-specific Rho1 knockdown (*moody>Rho1RNAi*) animals in single confocal sections of dye injected third instar larvae (C), and SJ morphology using the NrgGFP marker (D), with width of SJ belt highlighted by arrows. (E) Quantification of the dye penetration assay from (C). Columns represent intensity of dye penetration as measured by mean pixel intensity and normalized to WT mean (see Experimental Procedures), ±SEM, n=13-19. (F) Quantification of the mean diameter of SJ belts from (D), ±SEM, n= 8-13. In (A), (B) and (E) the percentage of animals showing dye penetration is indicated at the bottom of each column. Asterisks in (A), (B), (E) and (F) indicate significance levels of comparisons against either WT in (A), (E) and (F) or *PkaC1*^*B*3^ group in (B) based on one-way ANOVA with Dunnett’s multiple comparisons test in (A) and (B) or Welch’s ANOVA with Dunnett’s T3 multiple comparison test (E) and (F), n.s. p>0.05; *p<0.05; **p<0.01, ***p<0.001.

PKA is also known to phosphorylate and inhibit the small GTPase Rho1, which reduces the activity of its effector Rho kinase (ROK), ultimately resulting in decreased MLC phosphorylation and actomyosin contractility (Dong et al., 1998; Garcia et al., 1999; Howe, 2004; Lang et al., 1996; Tang et al., 2019; Xu and Myat, 2012). Moreover, RhoA activity has been shown to drive actin polymerization at the protrusion of migrating cells (Machacek et al., 2009), and a PKA-RhoA signaling has been suggested to act as a protrusion-retraction pacemaker at the leading edge of the migraing cells (Tkachenko et al., 2011). To check if Rho1 is required for BBB function, we determined the BBB permeability in the late embryo and third instar larval stages. Two loss-of-function alleles, the hypomorphic allele *Rho1*^*1B*^ (Magie and Parkhurst, 2005) and the null allele *Rho1*^*E*.*3*.*10*^ showed dye penetration defects as homozygous zygotic mutant embryos, with the null allele showing a particularly pronounced effect (Figure 6B and S5). At the larval stage, the SPG-specific Rho1 knockdown (*moody>Rho1RNAi*) resulted in strong dye penetration into the nerve cord (Figure 6C and 6E). These results suggest that Rho1 is required for the formation and continued growth of the BBB. We again asked whether PKA and Rho1 function in the same pathway and performed dominant genetic interaction experiments using a sensitized genetic background. The embryonic dye penetration defects of PkaC1 heterozygous mutants (*PkaC1*^*B3*^) were significantly reduced by removing one genomic copy of the Rho1 null allele (*Rho1*^*E*.*3*.*10*^), but not by removing one copy of the hypomorphic allele *Rho1*^*1B*^ (Figure 6B). These findings suggest that Rho1 is a PKA target in BBB regulation. Collectively, our results indicate that PKA suppresses actomyosin contractility in a two-pronged fashion, by negatively regulating both MLCK and Rho1.

## Discussion

Previous studies implicated a novel GPCR signaling pathway in the formation of the *Drosophila* BBB in late embryos (Bainton et al., 2005; Schwabe et al., 2005). This work also revealed that besides the GPCR Moody, two heterotrimeric G proteins (Gαiβγ, Gαoβγ) and the RGS Loco participate in this pathway. Here we provide a comprehensive molecular and cellular analysis of the events downstream of G protein signaling using a candidate gene screening approach. We present new, more sensitive methods for phenotypic characterization, and extended the analysis beyond the embryo into larval stages. This work identifies PKA, together with some of its targets, as crucial antagonistic effectors in the continued cell growth of SPG and maintenance of the BBB sealing capacity. This role is critical to ensure proper neuronal function during BBB formation and maturation.

Multiple lines of evidence demonstrate a role of PKA for proper sealing of the BBB: loss of PKA activity lead to to BBB permeability defects, irregular growth of SPG during epithelium formation, reduced membrane overlap and a narrower SJ belt at SPG cell-cell contacts. The role of PKA as an effector of the Moody signaling pathway is further supported by dominant genetic interaction experiments, which show that the dye penetration phenotype of *PkaC1* heterozygous mutant embryos was partially rescued by removing one genomic copy of *G*β*13F* or *loco*. Moreover, the analysis of the larval phenotype with live SJ and cytoskeleton markers shows that PKA gain-of-function behaved similarly to Moody loss-of-function. Conversely, PKA loss-of-function resembled the overexpression of GαoGTP, which mimics Moody gain-of-function signaling.

Our results from modulating PKA activity suggest that the total cell contact and SJ areas are a monotonic function of PKA activity: low levels of activity cause narrow contacts, and high levels give rise to broad contacts. Moreover, the analysis of various cellular markers (actin, microtubules, SJs, vesicles) indicates that the circumferential cytoskeleton and delivery of SJ components respond proportionately to PKA activity. This, in turn, promotes the increase in cell contact and junction areas coordinately at the lateral side of SPG. Our experiments demonstrate that the modulation of the SPG membrane overlap by PKA proceeds, at least in part, through the regulation of actomyosin contractility, and that this involves the phosphorylation targets MLCK and Rho1. This suggests that crucial characteristics of PKA signaling are conserved across eukaryotic organisms (Bauman et al., 2004; Marks and Kalderon, 2011; Park et al., 2000; Taylor et al., 1990; Tkachenko et al., 2011; Walker et al., 2013).

At the ultrastructural level, our ssTEM analysis of the larval SPG epithelium clarifies the relationship between the inter-cell membrane overlaps and SJ organization and function. Across different PKA activity levels, the ratio of septate junction areas to the total cell contact area remained constant at about 30%. This proportionality suggests a mechanism that couples cell contact with SJ formation. In this process the primary job of Moody/PKA appears to be the control of membrane overlap. This is consistent with the results of a temporal analysis of epithelium formation and SJ insertion in late embryos of WT and Moody pathway mutants, which shows that membrane contact precedes and is necessary for the appearance of SJs (Schwabe et al., 2017). The finding that the surface area that SJs occupy did not exceed a specific ratio, irrespective of the absolute area of cell contact, suggests an intrinsic, possibly steric limitation in how much junction can be fitted into a given cell contact space. While most phenotypic effects are indeed a monotonic function of Moody and PKA activity, the discontinuity and shortening of individual SJ strands is not. It occured with both increased and decreased signaling and appears to cause the leakiness of the BBB in both conditions. Our ssTEM-based 3D reconstruction thus demonstrates that the total area covered by SJs and the length of individual contiguous SJ segments are independent parameters. The latter appears to be critical for the paracellular seal, consistent with the idea that Moody plays a role in the formation of continuous SJ stands (Babatz et al., 2018).

The asymmetric localization of PKA that we observed sheds further light on the establishment and function of apical-basal polarity in the SPG epithelium. Prior to epithelium formation, contact with the basal lamina leads to the first sign of polarity (Schwabe et al., 2017). Moody becomes localized to the apical surface only after epithelial closure and SJ formation, suggesting that SJs are required as a diffusion barrier and that apical accumulation of Moody protein is the result of polarized exocytosis or endocytosis (Schwabe et al., 2017). Here, we now show that the intracellular protein PKA catalytic subunit-PkaC1 accumulates on the basal side of SPG, and that this polarized accumulation requires (apical) Moody activity. Such an asymmetric, activity-dependent localization has not previously been described for PKA in endothelium, and while the underlying molecular mechanism is unknown, the finding underscores that generating polarized activity along the apical-basal axis of the SPG is a key element of Moody pathway function.

An intriguing unresolved question is how increased SPG cell size and SJ length can keep up with the expanding brain without disrupting the BBB integrity during larva growth. We found that the SJ grows dramatically in length (0.57 ± 0.07 µm vs 2.16 ± 0.14 µm, about 3.7 fold) from the late embryo (Figure 1E) to 3rd instar larva (Figure 4G), which matches the increased cell size of SPG (about 4 fold) (Babatz et al., 2018; Unhavaithaya and Orr-Weaver, 2012). During the establishment of the SPG epithelium in the embryo, both increased and decreased Moody signaling resulted in asynchronous growth and cell contact formation along the circumference of SPG, which in turn led to irregular thickness of the SJ belt (Schwabe et al., 2017). Therefore, a similar relationship may exist during the continued growth of the SPG epithelium in larvae, with the loss of continuity of SJ segments in Moody/PKA mutants resulting from unsynchronized expansion of the cell contact area and an ensuing erratic insertion of SJ components. Since SJs form relatively static complexes, any irregularities in their delivery and insertion may linger for extended periods of time (Babatz et al., 2018; Deligiannaki et al., 2015; Oshima and Fehon, 2011). The idea that shortened SJ segments are a secondary consequence of unsynchronized cell growth is strongly supported by our finding that disruption of actomyosin contractility in MLCK and Rho1 mutants compromises BBB permeability.

Collectively, our data suggest the following model: polarized Moody/PKA signaling controls the cell growth and maintains BBB integrity during the continuous morphogenesis of the SPG secondary epithelium. On the apical side, Moody activity represses PKA activity (restricting local cAMP level within the apial-basal axis in SPG) and thereby promotes actomyosin contractility. On the basal side, which first adheres to the basal lamina and later to the PNG sheath, PKA activity suppresses actomyosin contractility via MLCK and Rho1 phosphorylation and repression (Figure 7). Throughout development, the SPG grow continuously while extending both their cell surface and expanding their cell contacts. Our data suggest that the membrane extension occurs on the basolateral surface through insertion of plasma membrane and cell-adhesive proteins, with similar behavior in epithelial cell, but regulated by a distinct polarized Moody/PKA signaling in SPG (Wojtal et al., 2008). In analogy to motile cells, the basal side of the SPG would thus act as the ‘leading edge’ of the cell, while the apical side functions as the ‘contractile rear’ (Nelson, 2009). According to this model, Moody/Rho1 regulate actomyosin to generate the contractile forces at the apical side to driving membrane contraction, which directs the basolateral insertion of new membrane material and SJs. In this way, differential contractility and membrane insertion act as a conveyor belt to move new formed membrane contacts and SJ from the basolateral to apical side. Loss of Moody signaling leads to symmetrical localization of PKA and to larger cell contact areas between SPG due to diminished apical constriction. Conversely, loss of PKA causes smaller cell contact areas due to increased basal constriction.

**Figure 7.**
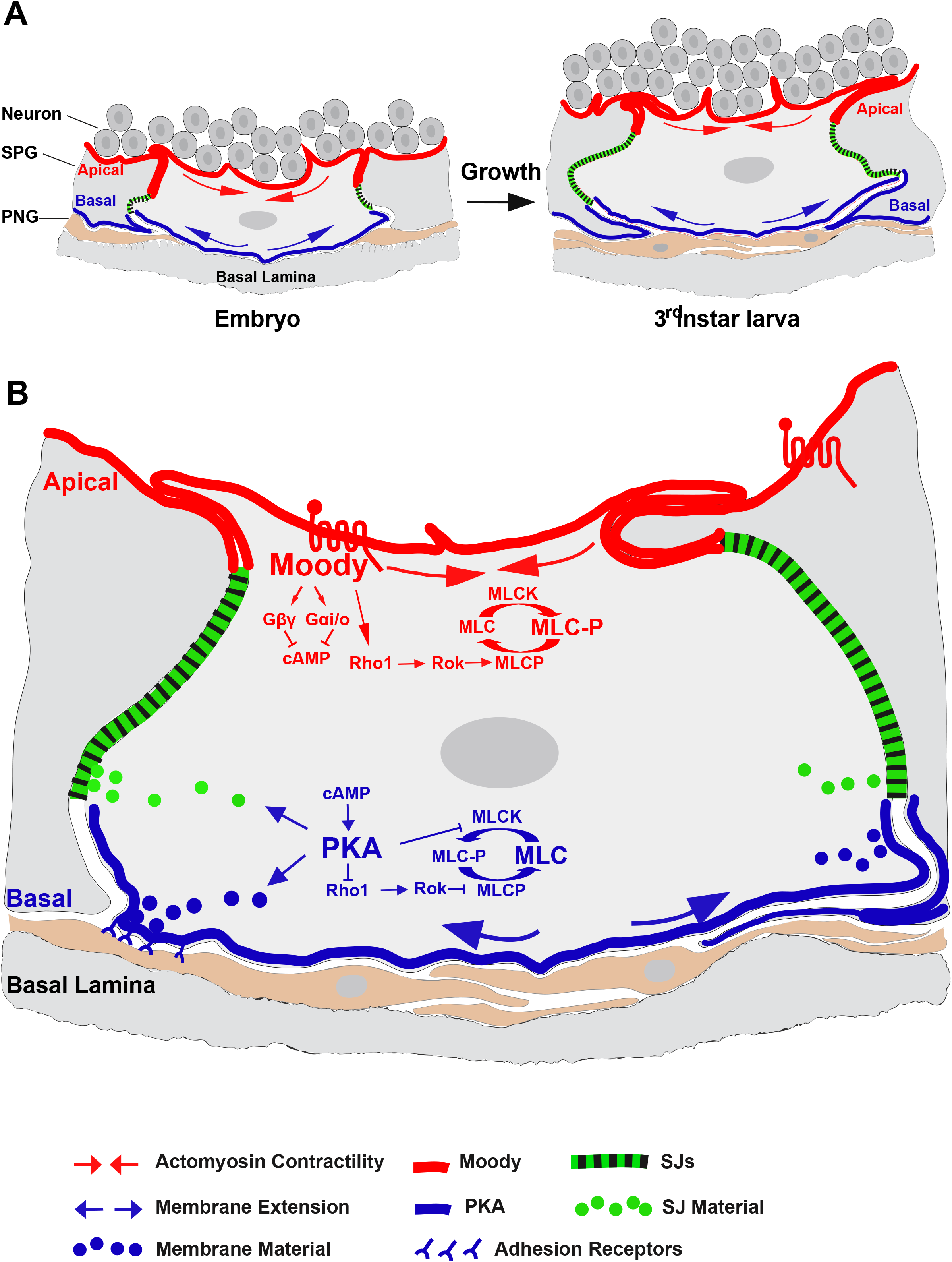
Model of Moody/PKA signaling in the glial BBB. Schematic depicting polarized Moody/PKA signaling along the apical-basal axis and its cellular function in controlling SPG continued cell growth and BBB integrity by differentially regulating actomyosin contractility and SJ organization spatiotemporally. For detailed description see Discussion.

Our results may have important implications for the development and maintenance of the BBB in vertebrates. The vertebrates BBB consistes of a secondary epithelium with interdigitations similar to the ones between the *Drosophila* SPG (Chow and Gu, 2015; Cong and Kong, 2020; Hindle and Bainton, 2014; Reinhold and Rittner, 2017). While the sealing is performed by tight junctions, it will be interesting to investigate whether there are similarities in the underlying molecular and cellular mechanisms that mediate BBB function (Artiushin et al., 2018; Cong and Kong, 2020; Sugimoto et al., 2020).

## Experimental Procedures

### Fly Strains and Constructs

The following fly strains were obtained from published sources: *PkaC1*^*H2*^ (BDSC Cat# 4101, RRID:BDSC_4101); *PkaC1*^*B3*^; *PkaC1*^*A13*^; *UAS*mPkaC1\**(mC*)*(D. Kalderon); *moodyGAL4* (T. Schwabe); *repoGAL4* (V. Auld); *Nrg*^*G305*^ (*NrgGFP*; W.Chia); *UASGFPMoesin* (D.Kiehart); *UAS*mRFPMoesin (T. Schwabe); *Gβ13F*^*Δ1-96A*^ (F. Matsuzaki); *UAStauGFP* (M. Krasnow); *UAS*Gαo*GTP* (A. Tomlinson), *loco*^*Δ13*^ (C. Klämbt); *moody*^*Δ17*^(R. Bainton); *moody-RNAi* (R. Bainton); *UASnucmCherry* (T. Schwabe); *UASGFPEB1* (D. Brunner); *UASGFPNod, UASGFPRho, UASactinGFP, UASRab4RFP, Rho*^*72R*^, *Rho*^*1B*^, *MLCK*^*02860*^, *MLCK*^*C234*^, *tubGAL80*^*ts*^ (Bloomington Stock Center); *PkaC1*^*KK108966*^, *Rho1*^*KK108182*^, *Tau*^*GD8682*^(VDRC). For live genotyping, mutant and transgenic lines were balanced (*Kr::GFP*) (Casso et al., 1999) or positively marked using *nrgNrgGFP*. Temperature-sensitive control of gene expression in SPG is achieved by using a *tubGAL80ts; moodyGAL4* driver. All strains were raised at 25°C. except for *tubGAL80ts; moodyGAL4* crosses, which were raised at 18°C until 1 day after eclosion and then shifted to 29°.

### Live Imaging

Dissected third-instar larval cephalic complexes were mounted in PBS and imaged directly. All confocal images were acquired using a Zeiss LSM 510 or 710 system. Stacks of 20-40 0.5 µm confocal sections were generated; image analysis was performed using Zeiss LSM 510, Image J (NIH) or Imaris 4.0 (Bitplane) software. The results for each section were assembled as a separate channel of the stack. Time-lapse recordings were carried out on 12h AEL embryos raised at 20°C using an inverted Zeiss LSM 510 confocal microscope. To increase signal strength, the pinhole was opened to 1.3 (z-section thickness 0.6 µm), and z-stacks of 12 sections were acquired once per minute. To adjust for focus drift, which is mainly caused by rotation of the embryo, the z-stack coordinates were adjusted at various time-points without disrupting the continuity of the movie. Between 5 and 7 movies were captured per genotype, each 80-110 min in duration.

### Immunohistochemistry

Immunohistochemistry was performed following standard procedures (Bainton et al., 2005; Schwabe et al., 2005). The antibodies used in the study were: rabbit α-PkaC1 (1:400, Pka-C1, RRID:AB_2568479) (Lane and Kalderon, 1993), mouse α-PkaC1 (1:100, BD), mouse α-REPO (1:10, Developmental Studies Hybridoma Bank), mouse α-GFP (1:100, Molecular Probes), mouse α-Mega (1:100, R. Schuh), guinea pig α-dContactin (1:1000, M. Bhat), rabbit α-RFP (1:100, US Biological). Fluorescent secondary antibodies were coupled to Cy3 (1:500, Jackson), Alexa Fluor 488 or Alexa Fluor 633 (1:500, Molecular Probes). Rat α-Moody β was generated in the lab (1:500).

### Image analysis

The width of the SJ belt was extracted from Maximum Intensity Projections (MIP) along the z-axis of 3D confocal stacks of the nervous system. Specifically, we used Imaris 4.0 (Bitplane) to perform 2D segmentation of the GFP-marked SJs. For each of the markers, an optimal threshold for the pixel intensity was chosen by fitting the obtained segmented pattern with the raw fluorescence signal. To evaluate the average thickness of the SJs, we split the SJ segments into sections of 3-4µm in length. An approximation of the diameters of the single sections was then obtained by extracting their ellipticity parameters along the axis perpendicular to their main axis. A mean diameter of the SJ was calculated by averaging over the diameters of all single sections.

### Dye-penetration assay in embryo, third instar larva, and adult flies

The dye penetration assay in embryos was performed as described (Schwabe et al., 2005). For the dye penetration assay in third instar larvae, a fluorescent dye (Texas red-coupled dextran, 10 kDa, 10mg/ml, Molecular Probes) was injected into the body cavity of third instar larva. After 2.5 h, the cephalic complex was dissected, and the dye penetrated into the nerve cord was analyzed using Zeiss LSM710 confocal microscopy. Dye penetration was quantified by calculating the percentage of larva showing dye penetration and by measuring the mean pixel intensity within a representative window of the ventral portion of the nerve cord using Fiji software, and normalized by dividing by the mean of the WT control group. To assess the significance of effects for the embryonic and larval dye penetration assays either ordinary or Welch’s ANOVA was performed, with Dunnett’s/Dunnett’s T3 or Tukey’s multiple comparisons test.

The dye penetration assay in adult flies was performed as described in Bainton et al. (2005) with some modifications. Briefly, adult flies were hemolymph injected with 10mg/ml 10kDa Texas red-coupled dextran. After 2h, the injected flies were decapitated and their heads were mounted in a fluorinated grease covered glass slides with two compound eyes on the side (the proboscis facing up). Images were acquired on a Zeiss LSM710 confocal microscope at 200-300 μm depths from the eye surface with a Plan Fluor 10xw objective. Dye penetration was quantified by measuring the mean pixel intensities within a representative window of the central region of retina (n=18-30) of maximum-intensity Z projection of each image stack (z-section thickness 0.6 µm) by Fiji software, and normalized by dividing by the WT control. Statistical significance was assessed using the two-tailed t-test.

### Transmission Electron Microscopy (TEM)

Late Stage 17 (22-23 hr AEL) embryos were processed by high pressure freezing in 20% BSA, freeze-substituted with 2% OsO4, 1% glutaraldehyde and 0.2% uranyl acetate in acetone (90%), dH2O (5%), methanol (5%) over 3 days (−90°C to 0°C), washed with acetone on ice, replaced with ethanol, infiltrated and embedded in Spurr’s resin, sectioned at 80 nm and stained with 2% uranyl acetate and 1% lead citrate for 5 min each. Sections were examined with a FEI TECNAI G2 Spirit BioTwin TEM with a Gatan 4K x 4K digital camera. For quantification, random images were shot, and the length of visible SJ membrane stretches in each image was measured using Fiji software. Statistics were calculated using the two-tailed Student’s t-test.

### Serial-section transmission electron microscopy (ssTEM)

Freshly dissected third instar larval CNSs were fixed in 2% glutaraldehyde and 2% OsO_4_in 0.12 M sodium cacodylate (pH 7.4) by microwave (Ted Pella, BioWave Pro MW) as follows: 30’’ at 300W, 60’’ OFF, 30’’ at 350W; 60’’ OFF, 30’’ at 400W. The samples were then rinsed 2×5’ with cold 0.12 M sodium cacodylate buffer; post-fixed with 1% OsO_4_in 0.12 M sodium cacodylate buffer (pH 7.4) on an ice bath by microwave as follows: 30’’ at 350W, 60’’ OFF, 30’’ at 375W, 60’’ OFF, 30’’ at 400W; rinsed 2×5’ with 0.12 M sodium cacodylate buffer at RT; 2×5’ with distilled water at RT; stained in 1% uranyl acetate overnight in 4°C; rinsed 6×5’ with distilled water; dehydrated with ethanol followed by propylene oxide (15’); infiltrated and embedded in Eponate 12 with 48h polymerization in a 65°C oven. 50 nm serial sections were cut on a Leica UC6 ultramicrotome and picked up with Synaptek slot grids on a carbon coated Pioloform film. Sections were post-stained with 1% uranyl acetate followed by Sato’s (1968) lead. The image acquisition of multiple sections (∼150 sections in each genotype) and large tissue areas were automatically captured with a Gatan 895 4K x 4K camera by a FEI Spirit TECNAI BioTWIN TEM using Leginon (Suloway et al., 2005). TrakEM2 software was used to montage, align images, trace and reconstruct 3D SJ structures between contacting SPG within and across serial sections. For quantification, random images were chosen, and the length of visible SJs stretches and membrane contacting area in each image were measured using Fiji. The statistical analysis was performed using Welch’s ANOVA with Dunnett’s T3 multiple comparisons test.

## Author contributions

X.L. and U.G. conceived and designed the study, X.L. and R.F. performed experiments, X.L., R.F., T.S., C.J., and U.G. analyzed and discussed data, H.S. provided laboratory resources and advice, X.L., U.G., and H.S. wrote the manuscript.

## Acknowledgements

We would like to thank D. Kalderon, R. Bainton, G. Beitel, V. Auld, M. Bhat, W. Chia, Y. Hiromi, M. Hortsch, B. Jones, D. Kiehart, C. Klämbt, J. Knoblich, M. Krasnow, M. Peifer, A. Tomlinson, R. Tsien, and the Developmental Studies Hybridoma Bank for providing us with fly strains, constructs, and antibodies. Special thanks go to U. Unnerstall, M. Deligiannaki, A. Casper, M. Schroeder,S. Axelrod, and E. Kurant for their helpful comments on the manuscript. We are grateful to all members of the Gaul and Steller labs for their continued support of this work.

## References

Artiushin, G., Zhang, S.L., Tricoire, H., and Sehgal, A. (2018). Endocytosis at the Drosophila blood-brain barrier as a function for sleep. Elife 7.

Babatz, F., Naffin, E., and Klambt, C. (2018). The Drosophila Blood-Brain Barrier Adapts to Cell Growth by Unfolding of Pre-existing Septate Junctions. Dev Cell 47, 697–710 e693.

Bainton, R.J., Tsai, L.T., Schwabe, T., DeSalvo, M., Gaul, U., and Heberlein, U. (2005). moody encodes two GPCRs that regulate cocaine behaviors and blood-brain barrier permeability in Drosophila. Cell 123, 145–156.

Banerjee, S., Sousa, A.D., and Bhat, M.A. (2006). Organization and function of septate junctions: an evolutionary perspective. Cell biochemistry and biophysics 46, 65–77.

Bauman, A.L., Goehring, A.S., and Scott, J.D. (2004). Orchestration of synaptic plasticity through AKAP signaling complexes. Neuropharmacology 46, 299–310.

Behr, M., Riedel, D., and Schuh, R. (2003). The claudin-like Megatrachea is essential in septate junctions for the epithelial barrier function in Drosophila. Developmental Cell 5, 611–620.

Cardona, A., Saalfeld, S., Schindelin, J., Arganda-Carreras, I., Preibisch, S., Longair, M., Tomancak, P., Hartenstein, V., and Douglas, R.J. (2012). TrakEM2 software for neural circuit reconstruction. PLoS One 7, e38011.

Carlson, S.D., Juang, J.L., Hilgers, S.L., and Garment, M.B. (2000). Blood barriers of the insect. Annual review of entomology 45, 151–174.

Casso, D., Ramirez-Weber, F.A., and Kornberg, T.B. (1999). GFP-tagged balancer chromosomes for Drosophila melanogaster. Mech Dev 88, 229–232.

Chen, X., and Ganetzky, B. (2012). A neuropeptide signaling pathway regulates synaptic growth in Drosophila. J Cell Biol 196, 529–543.

Chow, B.W., and Gu, C. (2015). The molecular constituents of the blood-brain barrier. Trends Neurosci 38, 598–608.

Clark, I.E., Jan, L.Y., and Jan, Y.N. (1997). Reciprocal localization of Nod and kinesin fusion proteins indicates microtubule polarity in the Drosophila oocyte, epithelium, neuron and muscle. Development 124, 461–470.

Cong, X., and Kong, W. (2020). Endothelial tight junctions and their regulatory signaling pathways in vascular homeostasis and disease. Cellular signalling 66, 109485.

Cui, W., Sproul, L.R., Gustafson, S.M., Matthies, H.J., Gilbert, S.P., and Hawley, R.S. (2005). Drosophila Nod protein binds preferentially to the plus ends of microtubules and promotes microtubule polymerization in vitro. Mol Biol Cell 16, 5400–5409.

Deligiannaki, M., Casper, A.L., Jung, C., and Gaul, U. (2015). Pasiflora proteins are novel core components of the septate junction. Development 142, 3046–3057.

Dong, J.M., Leung, T., Manser, E., and Lim, L. (1998). cA MP-induced Morphological Changes Are Counteracted by the Activated RhoA Small GTPase and the Rho Kinase ROKalpha. Journal of Biological Chemistry 273, 22554–22562.

Edwards, J.S., Swales, L.S., and Bate, M. (1993). The Differentiation between Neuroglia and Connective-Tissue Sheath in Insect Ganglia Revisited - the Neural Lamella and Perineurial Sheath-Cells Are Absent in a Mesodermless Mutant of Drosophila. J Comp Neurol 333, 301–308.

Faivre-Sarrailh, C., Banerjee, S., Li, J.J., Hortsch, M., Laval, M., and Bhat, M.A. (2004). Drosophila contactin, a homolog of vertebrate contactin, is required for septate junction organization and paracellular barrier function. Development 131, 4931–4942.

Fessler, L.I., Nelson, R.E., and Fessler, J.H. (1994). Drosophila extracellular matrix. Methods in enzymology 245, 271–294.

Garcia, J.G., Davis, H.W., and Patterson, C.E. (1995). Regulation of endothelial cell gap formation and barrier dysfunction: role of myosin light chain phosphorylation. Journal of cellular physiology 163, 510–522.

Garcia, J.G., Lazar, V., Gilbert-McClain, L.I., Gallagher, P.J., and Verin, A.D. (1997). Myosin light chain kinase in endothelium: molecular cloning and regulation. American journal of respiratory cell and molecular biology 16, 489–494.

Garcia, J.G., Verin, A.D., Schaphorst, K., Siddiqui, R., Patterson, C.E., Csortos, C., and Natarajan,V. (1999). Regulation of endothelial cell myosin light chain kinase by Rho, cortactin, and p60(src). Am J Physiol 276, L989–998.

Genova, J.L., and Fehon, R.G. (2003). Neuroglian, Gliotactin, and the Na+/K+ ATPase are essential for septate junction function in Drosophila. J Cell Biol 161, 979–989.

Granderath, S., Stollewerk, A., Greig, S., Goodman, C.S., O’Kane, C.J., and Klambt, C. (1999). loco encodes an RGS protein required for Drosophila glial differentiation. Development 126, 1781–1791.

Guan, Z., Buhl, L.K., Quinn, W.G., and Littleton, J.T. (2011). Altered gene regulation and synaptic morphology in Drosophila learning and memory mutants. Learning & memory 18, 191–206.

Halter, D.A., Urban, J., Rickert, C., Ner, S.S., Ito, K., Travers, A.A., and Technau, G.M. (1995). The Homeobox Gene Repo Is Required for the Differentiation and Maintenance of Glia Function in the Embryonic Nervous-System of Drosophila-Melanogaster. Development 121, 317–332.

Hartenstein, V. (2011). Morphological diversity and development of glia in Drosophila. Glia 59, 1237–1252.

Hijazi, A., Haenlin, M., Waltzer, L., and Roch, F. (2011). The Ly6 protein coiled is required for septate junction and blood brain barrier organisation in Drosophila. PLoS One 6, e17763.

Hijazi, A., Masson, W., Auge, B., Waltzer, L., Haenlin, M., and Roch, F. (2009). boudin is required for septate junction organisation in Drosophila and codes for a diffusible protein of the Ly6 superfamily. Development 136, 2199–2209.

Hindle, S.J., and Bainton, R.J. (2014). Barrier mechanisms in the Drosophila blood-brain barrier. Front Neurosci 8, 414.

Howe, A.K. (2004). Regulation of actin-based cell migration by cAMP/PKA. Biochim Biophys Acta 1692, 159–174.

Ito, K., Urban, J., and Technau, G.M. (1995). Distribution, classification, and development ofDrosophila glial cells in the late embryonic and early larval ventral nerve cord. Roux’s Arch Dev Biol 204, 284–307.

Izumi, Y., and Furuse, M. (2014). Molecular organization and function of invertebrate occluding junctions. Semin Cell Dev Biol 36, 186–193.

Jarecki, J., Johnson, E., and Krasnow, M.A. (1999). Oxygen regulation of airway branching in Drosophila is mediated by branchless FGF. Cell 99, 211–220.

Kalderon, D., and Rubin, G.M. (1988). Isolation and characterization of Drosophila cAMP-dependent protein kinase genes. Genes Dev 2, 1539–1556.

Lane, M.E., and Kalderon, D. (1993). Genetic investigation of cAMP-dependent protein kinase function in Drosophila development. Genes Dev 7, 1229–1243.

Lane, M.E., and Kalderon, D. (1994). RNA localization along the anteroposterior axis of the Drosophila oocyte requires PKA-mediated signal transduction to direct normal microtubule organization. Genes Dev 8, 2986–2995.

Lane, M.E., and Kalderon, D. (1995). Localization and functions of protein kinase A during Drosophila oogenesis. Mech Dev 49, 191–200.

Lang, P., Gesbert, F., espine-Carmagnat, M., Stancou, R., Pouchelet, M., and Bertoglio, J. (1996). Protein kinase A phosphorylation of RhoA mediates the morphological and functional effects of cyclic AMP in cytotoxic lymphocytes. EMBO J 15, 510–519.

Li, D., Liu, Y., Pei, C., Zhang, P., Pan, L., Xiao, J., Meng, S., Yuan, Z., and Bi, X. (2017). miR-285-Yki/Mask double-negative feedback loop mediates blood-brain barrier integrity in Drosophila. Proc Natl Acad Sci U S A 114, E2365–E2374.

Li, W., Ohlmeyer, J.T., Lane, M.E., and Kalderon, D. (1995). Function of protein kinase A in hedgehog signal transduction and Drosophila imaginal disc development. Cell 80, 553–562.

Li, W., Tully, T., and Kalderon, D. (1996). Effects of a conditional Drosophila PKA mutant on olfactory learning and memory. Learning & memory 2, 320–333.

Lim, C.J., Kain, K.H., Tkachenko, E., Goldfinger, L.E., Gutierrez, E., Allen, M.D., Groisman, A., Zhang, J., and Ginsberg, M.H. (2008). Integrin-mediated Protein Kinase A Activation at the Leading Edge of Migrating Cells. Molecular Biology of the Cell 19, 4930–4941.

Llimargas, M., Strigini, M., Katidou, M., Karagogeos, D., and Casanova, J. (2004). Lachesin is a component of a septate junction-based mechanism that controls tube size and epithelial integrity in the Drosophila tracheal system. Development 131, 181–190.

Machacek, M., Hodgson, L., Welch, C., Elliott, H., Pertz, O., Nalbant, P., Abell, A., Johnson, G.L., Hahn, K.M., and Danuser, G. (2009). Coordination of Rho GTPase activities during cell protrusion. Nature 461, 99–103.

Magie, C.R., and Parkhurst, S.M. (2005). Rho1 regulates signaling events required for proper Drosophila embryonic development. Dev Biol 278, 144–154.

Marks, S.A., and Kalderon, D. (2011). Regulation of mammalian Gli proteins by Costal 2 and PKA in Drosophila reveals Hedgehog pathway conservation. Development 138, 2533–2542.

Mayer, F., Mayer, N., Chinn, L., Pinsonneault, R.L., Kroetz, D., and Bainton, R.J. (2009). Evolutionary conservation of vertebrate blood-brain barrier chemoprotective mechanisms in Drosophila. The Journal of neuroscience: the official journal of the Society for Neuroscience 29, 3538–3550.

Nelson, W.J. (2009). Remodeling epithelial cell organization: transitions between front-rear and apical-basal polarity. Cold Spring Harbor perspectives in biology 1, a000513.

Olofsson, B., and Page, D.T. (2005). Condensation of the central nervous system in embryonic Drosophila is inhibited by blocking hemocyte migration or neural activity. Dev Biol 279, 233–243.

Oshima, K., and Fehon, R.G. (2011). Analysis of protein dynamics within the septate junction reveals a highly stable core protein complex that does not include the basolateral polarity protein Discs large. Journal of Cell Science 124, 2861–2871.

Park, S.K., Sedore, S.A., Cronmiller, C., and Hirsh, J. (2000). Type II cAMP-dependent protein kinase-deficient Drosophila are viable but show developmental, circadian, and drug response phenotypes. J Biol Chem 275, 20588–20596.

Petri, J., Syed, M.H., Rey, S., and Klambt, C. (2019). Non-Cell-Autonomous Function of the GPI-Anchored Protein Undicht during Septate Junction Assembly. Cell Rep 26, 1641–1653 e1644.

Reinhold, A.K., and Rittner, H.L. (2017). Barrier function in the peripheral and central nervous system-a review. Pflugers Arch 469, 123–134.

Renger, J.J., Ueda, A., Atwood, H.L., Govind, C.K., and Wu, C.F. (2000). Role of cAMP cascade in synaptic stability and plasticity: ultrastructural and physiological analyses of individual synaptic boutons in Drosophila memory mutants. The Journal of neuroscience: the official journal of the Society for Neuroscience 20, 3980–3992.

Rogers, S.L., Wiedemann, U., Hacker, U., Turck, C., and Vale, R.D. (2004). Drosophila RhoGEF2 associates with microtubule plus ends in an EB1-dependent manner. Curr Biol 14, 1827–1833.

Salzer, J.L., Brophy, P.J., and Peles, E. (2008). Molecular domains of myelinated axons in the peripheral nervous system. Glia 56, 1532–1540.

Schindelin, J., Arganda-Carreras, I., Frise, E., Kaynig, V., Longair, M., Pietzsch, T., Preibisch, S., Rueden, C., Saalfeld, S., Schmid, B., et al. (2012). Fiji: an open-source platform for biological-image analysis. Nature methods 9, 676–682.

Schwabe, T., Bainton, R.J., Fetter, R.D., Heberlein, U., and Gaul, U. (2005). GPCR signaling is required for blood-brain barrier formation in Drosophila. Cell 123, 133–144.

Schwabe, T., Li, X., and Gaul, U. (2017). Dynamic analysis of the mesenchymal-epithelial transition of blood-brain barrier forming glia in Drosophila. Biol Open 6, 232–243.

Shabb, J.B. (2001). Physiological substrates of cAMP-dependent protein kinase. Chemical reviews 101, 2381–2411.

Stork, T., Engelen, D., Krudewig, A., Silies, M., Bainton, R.J., and Klambt, C. (2008). Organization and function of the blood-brain barrier in Drosophila. The Journal of neuroscience: the official journal of the Society for Neuroscience 28, 587–597.

Sugimoto, K., Ichikawa-Tomikawa, N., Nishiura, K., Kunii, Y., Sano, Y., Shimizu, F., Kakita, A., Kanda, T., Imura, T., and Chiba, H. (2020). Serotonin/5-HT1A Signaling in the Neurovascular Unit Regulates Endothelial CLDN5 Expression. Int J Mol Sci 22.

Suloway, C., Pulokas, J., Fellmann, D., Cheng, A., Guerra, F., Quispe, J., Stagg, S., Potter, C.S., and Carragher, B. (2005). Automated molecular microscopy: the new Leginon system. Journal of structural biology 151, 41–60.

Syed, M.H., Krudewig, A., Engelen, D., Stork, T., and Klambt, C. (2011). The CD59 family member Leaky/Coiled is required for the establishment of the blood-brain barrier in Drosophila. The Journal of neuroscience: the official journal of the Society for Neuroscience 31, 7876–7885.

Tang, S.-t., Tang, H.-q., Su, H., Wang, Y., Zhou, Q., Zhang, Q., Wang, Y., and Zhu, H.-q. (2019). Glucagon-like peptide-1 attenuates endothelial barrier injury in diabetes via cAMP/PKA mediated down-regulation of MLC phosphorylation. Biomedicine & Pharmacotherapy 113, 108667.

Taylor, S.S., Buechler, J.A., and Yonemoto, W. (1990). cAMP-dependent protein kinase: framework for a diverse family of regulatory enzymes. Annual review of biochemistry 59, 971–1005.

Tempesta, C., Hijazi, A., Moussian, B., and Roch, F. (2017). Boudin trafficking reveals the dynamic internalisation of specific septate junction components in Drosophila. PLoS One 12, e0185897.

Tepass, U. (2012). The apical polarity protein network in Drosophila epithelial cells: regulation of polarity, junctions, morphogenesis, cell growth, and survival. Annual review of cell and developmental biology 28, 655–685.

Tepass, U., and Hartenstein, V. (1994). The development of cellular junctions in the Drosophila embryo. Dev Biol 161, 563–596.

Tepass, U., Tanentzapf, G., Ward, R., and Fehon, R. (2001). Epithelial cell polarity and cell junctions in Drosophila. Annual review of genetics 35, 747–784.

Tkachenko, E., Sabouri-Ghomi, M., Pertz, O., Kim, C., Gutierrez, E., Machacek, M., Groisman, A., Danuser, G., and Ginsberg, M.H. (2011). Protein kinase A governs a RhoA-RhoGDI protrusion-retraction pacemaker in migrating cells. Nat Cell Biol 13, 660–667.

Unhavaithaya, Y., and Orr-Weaver, T.L. (2012). Polyploidization of glia in neural development links tissue growth to blood-brain barrier integrity. Genes Dev 26, 31–36.

Vasin, A., Zueva, L., Torrez, C., Volfson, D., Troy Littleton, J., and Bykhovskaia, M. (2014). Synapsin regulates activity-dependent outgrowth of synaptic boutons at the Drosophila neuromuscular junction. Journal of Neuroscience 34, 10554–10563.

Verin, A.D., Gilbert-McClain, L.I., Patterson, C.E., and Garcia, J.G.N. (1998). Biochemical Regulation of the Nonmuscle Myosin Light Chain Kinase Isoform in Bovine Endothelium. American Journal of Respiratory Cell and Molecular Biology 19, 767–776.

Von Stetina, J.R., Frawley, L.E., Unhavaithaya, Y., and Orr-Weaver, T.L. (2018). Variant cell cycles regulated by Notch signaling control cell size and ensure a functional blood-brain barrier. Development 145.

Walker, J.A., Gouzi, J.Y., Long, J.B., Huang, S., Maher, R.C., Xia, H., Khalil, K., Ray, A., Van Vactor, D., Bernards, R., et al. (2013). Genetic and functional studies implicate synaptic overgrowth and ring gland cAMP/PKA signaling defects in the Drosophila melanogaster neurofibromatosis-1 growth deficiency. PLoS genetics 9, e1003958.

Wojtal, K.A., Hoekstra, D., and van Ijzendoorn, S.C. (2008). cAMP-dependent protein kinase A and the dynamics of epithelial cell surface domains: moving membranes to keep in shape. BioEssays: news and reviews in molecular, cellular and developmental biology 30, 146–155.

Wu, V.M., Schulte, J., Hirschi, A., Tepass, U., and Beitel, G.J. (2004). Sinuous is a Drosophila claudin required for septate junction organization and epithelial tube size control. J Cell Biol 164, 313–323.

Xu, N., and Myat, M.M. (2012). Coordinated control of lumen size and collective migration in the salivary gland. Fly 6, 142–146.

Yi, P., Johnson, A.N., Han, Z., Wu, J., and Olson, E.N. (2008). Heterotrimeric G proteins regulate a noncanonical function of septate junction proteins to maintain cardiac integrity in Drosophila. Dev Cell 15, 704–713.

Zhang, J., Schulze, K.L., Hiesinger, P.R., Suyama, K., Wang, S., Fish, M., Acar, M., Hoskins, R.A., Bellen, H.J., and Scott, M.P. (2007). Thirty-one flavors of Drosophila rab proteins. Genetics 176, 1307–1322.

Zhou, Q., Apionishev, S., and Kalderon, D. (2006). The contributions of protein kinase A and smoothened phosphorylation to hedgehog signal transduction in Drosophila melanogaster. Genetics 173, 2049–2062.

