## Supplemental S1-S5 for "The cAMP effector PKA mediates Moody GPCR signaling in *Drosophila* blood-brain barrier formation and maturation"

Movie S1. related to Fig. 1F

Movie S2. related to Fig. 1F

Figure S1. related to Fig. 1G.

Figure S2. related to Fig. 2A.

Figure S3. related to Fig. 2B. and 2C.

Figure S4. related to Fig. 4A-E and Fig 7.

Figure S5. related to Fig. 6A.

Figure S1. related to Fig. 1G.

**Figure S1. Genetic interactions between PKA and different Moody pathway components.**

Quantitative analysis of dye penetration shows that the mild BBB defect of *PkaC1*$^{B3}$ heterozygotes can be partially rescued by removing one copy of *Gβ13F* as well as one copy of *loco.* The removal of one copy of other Moody pathway components, such as *moody*, *Gαo*, and *Gαi*, results in no or weaker, non-significant rescue, suggesting that *Gβ13F* is more dosage sensitive. Asterisks indicate statistical significance levels as assessed by one-way ANOVA with Dunnett's multiple comparisons test, n.s. $p > 0.05$; *$p<0.05$; **$p<0.01$; ***$p<0.001$.

Figure S1. related to Fig. 1G.

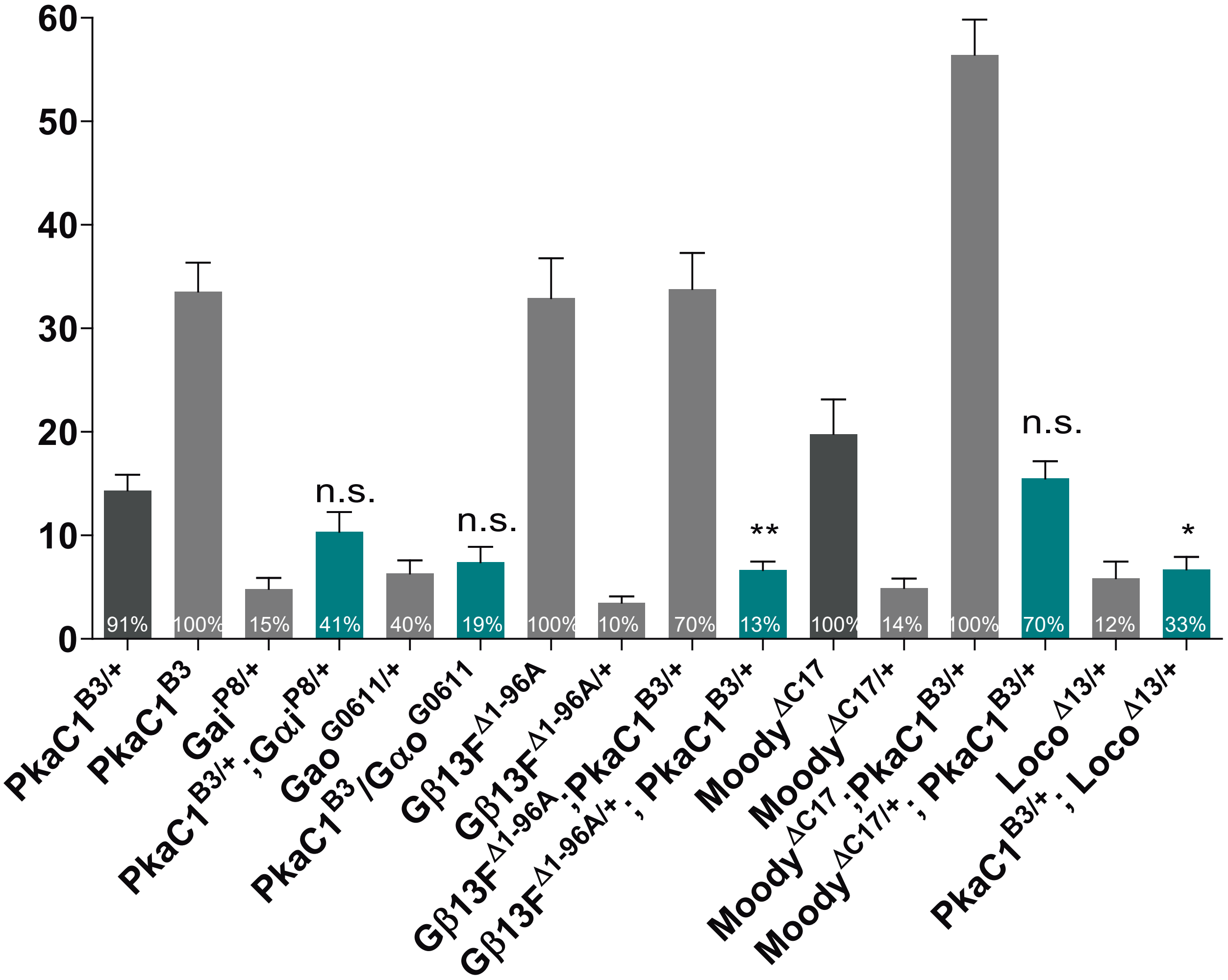

Figure S2. related to Fig. 2A.

**Figure S2. Dye penetration defects for RNAi vs. genomic mutants in the embryo.** Graph shows quantification of penetration of fluorescent dye into the CNS in WT, *loco* and *moody* null alleles, as well as *loco* or *moody*RNAi expressed under *repo*- or *moody-Gal4*; 22h embryos. While *moodyRNAi* expressed using *moody-Gal4* results in measurable dye penetration, the defect is much milder than in the mutant. Asterisks indicate statistical significance levels as assessed by one-way ANOVA with Dunnett's multiple comparisons test n.s. p> 0.05; *p<0.05; **p<0.01; ***p<0.001.

Figure S2. related to Fig. 2A.

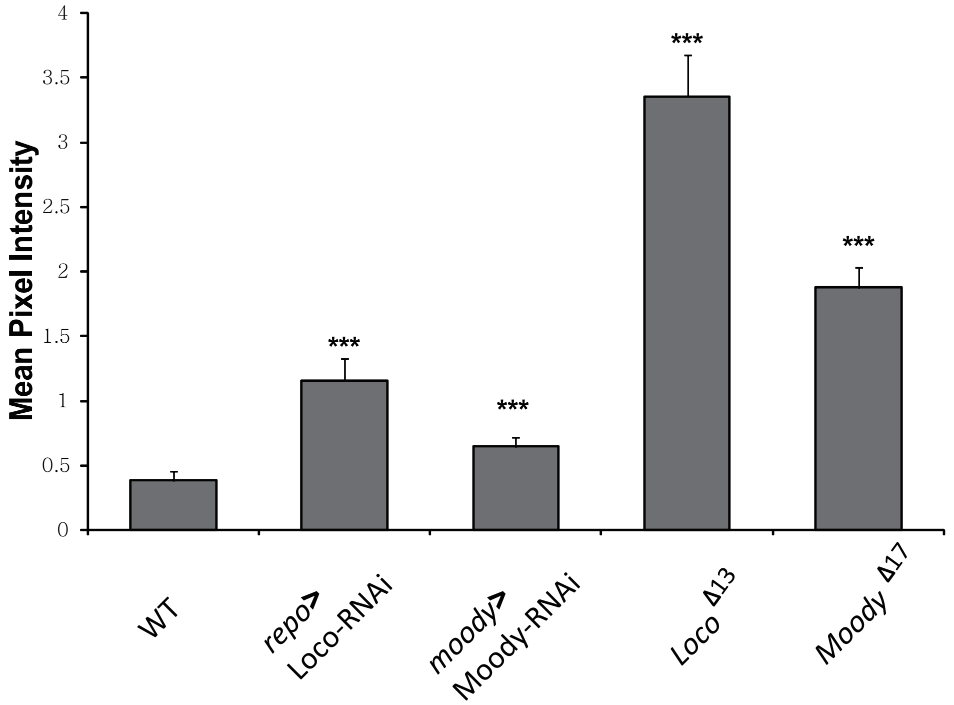

Figure S3. related to Fig. 2B. and 2C.

**Figure S3. Morphology of SPG SJ belts at different PKA activity levels**, as visualized by SJ markers *LacGFP* (A), *Mega* (B), and *Nrx-IVGFP* (C). (D) Quantification of the diameter of SJ belts under different PKA activity levels, using different SJ markers. All groups in PKA overactivity and one group of *LacGFP* labeled SJ belt in PKA underactivity are significantly different from WT, as assessed by Welch's ANOVA with Dunnett's T3 multiple comparisons test, n.s. $p>0.05$; $^{*}p<0.05$; $^{**}p<0.01$, $^{***}p<0.001$.

Figure S3. related to Fig. 2B. and 2C.

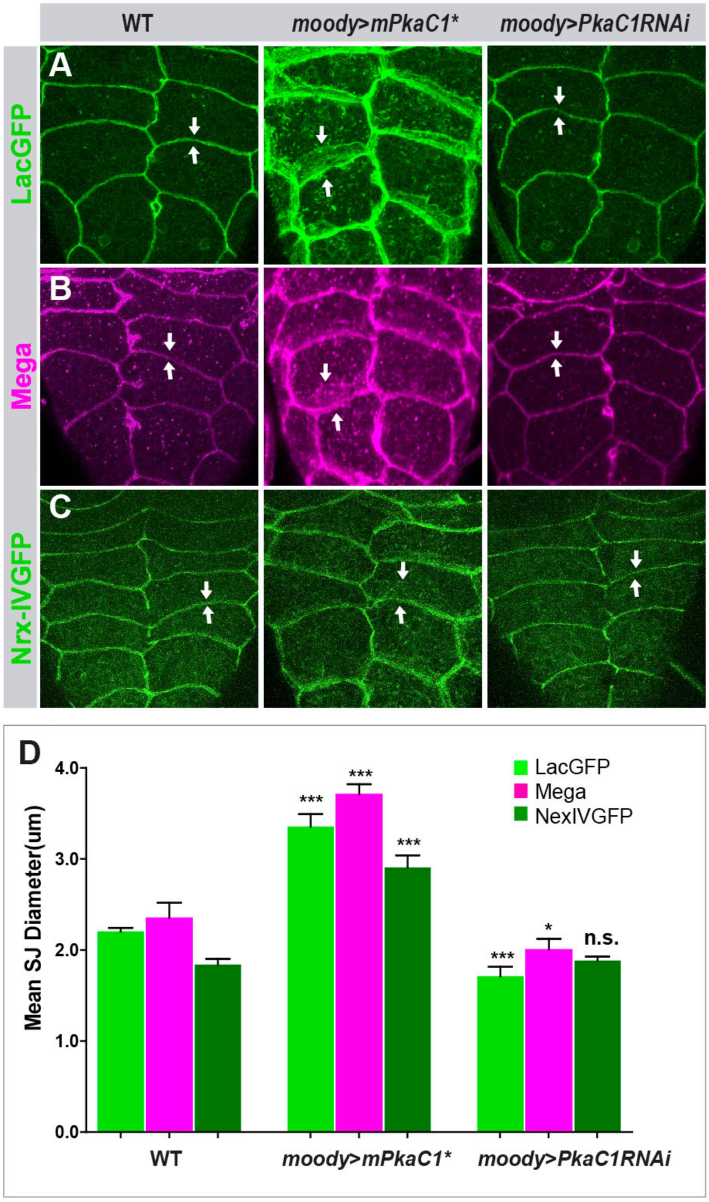

Figure S4. related to Fig. 4A-E.

**Figure S4. Morphology of apical membrane protrusions of SPG at different PKA activity levels, as revealed by ssTEM.** (A) Representative sections of the apical membrane protrusion of SPG ultrastructure under different PKA levels, marked by yellow arrows. SPG1, its neighbor SPG2, and their shared SJs are colored in red, magenta and green, respectively. (B) Quantification of the mean length of individual apical protrusion under different PKA activity levels, measured in random nerve cord sections, ±SEM, n=14-35. Compared to WT, the mean length of apical membrane protrusion of SPG is significantly longer under PKA overactivity and significantly shorter under PKA underactivity. Statistical significance of comparisons was assessed using Welch’s ANOVA with Dunnett's T3 multiple comparisons test, n.s. $p>0.05$; *$p<0.05$; **$p<0.01$, ***$p<0.001$.

Figure S4. related to Fig. 4A-E. and Fig. 7.

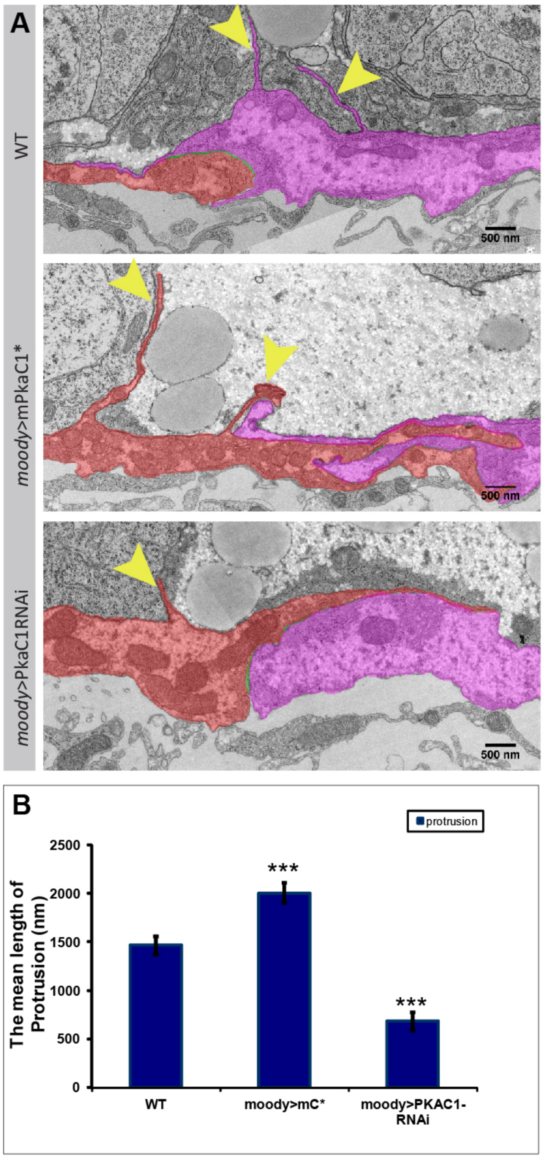

Figure S5. related to Fig. 6A.

**Figure S5. MLCK and Rho1 are required for BBB integrity in embryos.** Single confocal section of dye injected *MLCK*$^{C234}$ zygotic mutant (*MLCK*$^{C234}$), and Rho1 zygotic null mutant (*Rho1*$^{E3.10)}$ embryos, with dye penetrating into the nerve cord (yellow arrow).

Figure S5. related to Fig. 6A.

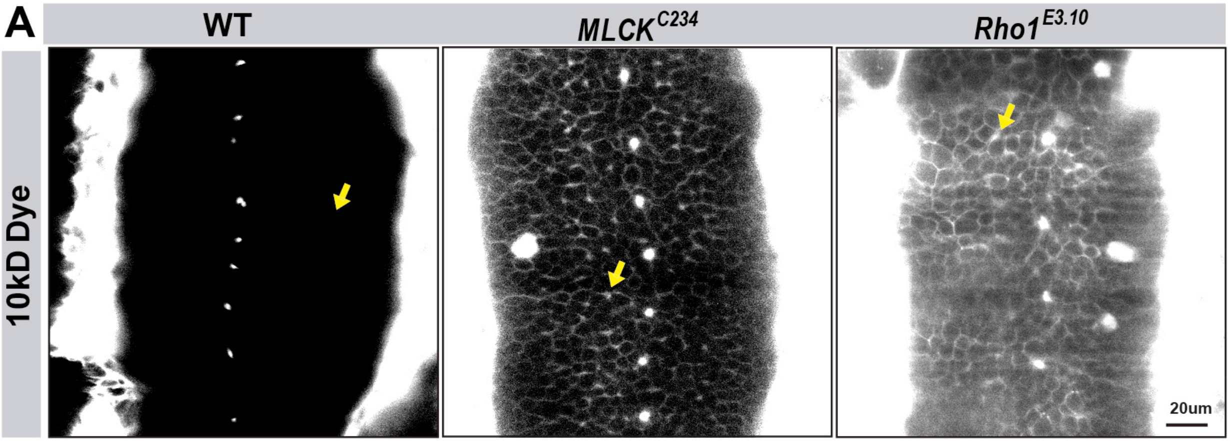

**Movie S1. SPG epithelium formation in WT embryos.** The SPG (labeled by the membrane marker *GapGFP* and the actin marker *MoesinGFP*, driven by the pan-glial driver *repoGAL4*) are fairly uniform in size and cell shape, and their spreading and cell-cell contact formation are highly synchronized. The glial sheet is closed by 15.5 h of development.

**Movie S2. SPG epithelium formation in PKA zygotic null mutant embryos.** The SPG of *PkaC1*$^{B3}$ zygotic mutants show variable size and cell shape, and their spreading and contact formation are poorly synchronized, resulting in patchy cell-cell contacts with gaps of different sizes. Moreover, the complete closure of the SPG epithelium is delayed compared to WT.
